# Hippocampal and striatal responses during motor learning are modulated by prefrontal cortex stimulation

**DOI:** 10.1101/2020.06.05.136531

**Authors:** Mareike A. Gann, Bradley R. King, Nina Dolfen, Menno P. Veldman, Kimberly L. Chan, Nicolaas A. J. Puts, Richard A. E. Edden, Marco Davare, Stephan P. Swinnen, Dante Mantini, Edwin M. Robertson, Geneviève Albouy

## Abstract

While it is widely accepted that motor sequence learning (MSL) is supported by a prefrontal-mediated interaction between hippocampal and striatal networks, it remains unknown whether the functional responses of these networks can be modulated in humans with targeted experimental interventions. The present proof-of-concept study employed a comprehensive multimodal neuroimaging approach, including functional magnetic resonance (MR) imaging and MR spectroscopy, to investigate whether individually-tailored theta-burst stimulation of the dorsolateral prefrontal cortex can modulate responses in the hippocampus and striatum during motor learning. Our results indicate that stimulation influenced task-related *connectivity* patterns within hippocampo-frontal and striatal networks. Stimulation also altered the relationship between the levels of gamma-aminobutyric acid (GABA) in the stimulated prefrontal cortex and learning-related changes in both *activity* and *connectivity* in fronto-striato-hippocampal networks. This study provides the first experimental evidence that brain stimulation can alter motor learning-related functional responses in the striatum and hippocampus.

## Introduction

The acquisition of new motor skills has been extensively studied using motor sequence learning (MSL) tasks during which participants integrate a series of movements into a temporally coherent structure. The neural responses underlying the initial learning phase have been thoroughly investigated and various models propose that MSL is supported by cortico-cerebellar, -striatal and -hippocampal networks (Albouy, King, Maquet, & Doyon, 2013; Doyon et al., 2009; Penhune & Steele, 2012). While cortico-striatal circuits have been described to support the development of a motoric representation of the sequence through practice, cortico-hippocampal networks are thought to promote the building of a spatial map of the sequence supporting a more abstract representation of the motor skill (Albouy et al., 2015; Albouy, King, et al., 2013).

Interestingly, the brain systems described above present different dynamical patterns of activity during the learning process (Albouy, King, et al., 2013). Whereas activity in hippocampo-fronto-parietal networks, which form loops with associative regions of the striatum and the cerebellum, decreases as a function of learning, activity in sensorimotor circuits, including the sensorimotor parts of the striatum, the cerebellum and motor cortical areas, increases with learning (Albouy, King, et al., 2013; Albouy et al., 2008, 2012; Doyon, Gabitov, Vahdat, Lungu, & Boutin, 2018; Hikosaka, Nakamura, Sakai, & Nakahara, 2002). Importantly, functional connectivity between these networks reveals a competitive interaction pattern during this initial learning stage (Albouy, King, et al., 2013; Albouy, Sterpenich, et al., 2013). Crucial to the present study, this interaction between hippocampal and striatal systems is thought to be orchestrated by the dorsolateral prefrontal cortex (DLPFC) (Albouy, King, et al., 2013; Albouy et al., 2012; Albouy, Sterpenich, et al., 2013; Freedberg, Toader, Wassermann, & Voss, 2020).

As the hippocampal and striatal neural signatures described above are thought to support motor memory acquisition and also predict successful motor memory retention (Albouy et al., 2008; Albouy, Sterpenich, et al., 2013; Steele & Penhune, 2010), investigating whether the amplitude and the dynamics of these learning-related brain responses can be altered by experimental interventions is of the utmost importance. One experimental approach that has shown promise to modulate neural responses in the striatum and hippocampus is the application of non-invasive brain stimulation to cortical regions that are functionally connected to these deep areas. For example, it has been shown that the application of repetitive transcranial magnetic stimulation (TMS) to the DLPFC (Bilek et al., 2013) or the parietal cortex (Freedberg et al., 2019; Wang et al., 2014) can alter the functional connectivity between the targeted cortical area and the hippocampus which, in turn, influences performance on working and associative memory tasks (Bilek et al., 2013; Wang et al., 2014). In addition, prefrontal TMS has been shown to influence striatal activity and connectivity at rest (Alkhasli, Sakreida, Mottaghy, & Binkofski, 2019; Esslinger et al., 2014; Hanlon, Dowdle, Moss, Canterberry, & George, 2016; van der Werf, Sanz-Arigita, Menning, & van den Heuvel, 2010) as well as during reward processing (Van Holstein, Froböse, O’Shea, Aarts, & Cools, 2018) and probabilistic learning (Ott, Ullsperger, Jocham, Neumann, & Klein, 2011). Based on the aforementioned evidence that the DLPFC mediates the interaction between the striato- and hippocampo-cortical systems during initial MSL and that prefrontal stimulation can influence functional responses in these networks, the DLPFC is a promising cortical stimulation target in order to alter brain responses in motor learning-relevant networks.

The goal of the present proof-of-concept study was therefore to use an extensive and multimodal neuroimaging approach, including functional Magnetic Resonance Imaging (fMRI) and MR Spectroscopy (MRS), to test, for the first time, whether stimulation of the DLPFC can modulate motor-learning-related functional responses in cortico-striatal and cortico-hippocampal networks. Based on evidence that the neuromodulatory effects of TMS can be optimized by (1) defining stimulation targets via data-driven approaches and (2) tailoring the stimulation targeting procedures to each individual (Beynel et al., 2019; Fox, Buckner, White, Greicius, & Pascual-Leone, 2012; Fox, Halko, Eldaief, & Pascual-Leone, 2012; Sack et al., 2009), we used a two-step approach. First, we analyzed fMRI data from a sample of young healthy individuals (Experiment 1) to identify a cortical cluster functionally connected to both the striatum and hippocampus at rest. In a second step, we used the identified spatial location to guide an individualized TMS targeting procedure (Wang et al., 2014) on an independent sample of young healthy participants (Experiment 2). In the second experiment, theta-burst stimulation (TBS), a form of repetitive TMS, was applied to the identified prefrontal cortical target ***before*** participants were trained on a sequential serial reaction time task (SRTT, Nissen & Bullemer, 1987) or a control random condition (random SRTT). Specifically, we examined the effect of intermittent versus continuous TBS (i.e., iTBS and cTBS, respectively; Huang, Edwards, Rounis, Bhatia, & Rothwell, 2005) of the DLPFC on (1) task-related activity and connectivity patterns measured with fMRI during post-stimulation task practice and (2) DLPFC and hippocampal neurochemistry through the quantification of gamma-aminobutyric acid (GABA), the brain’s primary inhibitory neurotransmitter, pre- and post-intervention using MRS.

As stimulation-induced effects of TBS on neural excitability have been shown to be similar in the prefrontal cortex as in the primary motor cortex (M1; Chung et al., 2017), we hypothesized that facilitatory iTBS and inhibitory cTBS of the DLPFC would respectively strengthen and disrupt activity and connectivity in hippocampo-parieto-prefrontal networks during sequence learning as compared to random practice. Based on models suggesting that hippocampo-prefrontal networks exert control processes over sensorimotor-striato-cortical networks during MSL (Albouy, King, et al., 2013), we expected that facilitatory iTBS and inhibitory cTBS of the DLPFC would repress and facilitate, respectively, the development of striato-motor activity during sequence learning as compared to random practice. With respect to the effects on GABA, previous MRS studies have shown that M1 GABA levels can be altered by both M1 brain stimulation (Bachtiar et al., 2018; Bachtiar, Near, Johansen-Berg, & Stagg, 2015; Marjańska et al., 2013; Stagg, Bachtiar, & Johansen-Berg, 2011; Stagg, Best, et al., 2009; Stagg, Wylezinska, et al., 2009) and motor learning (Floyer-Lea, Wylezinska, Kincses, & Matthews, 2006; Kolasinski et al., 2018; Sampaio-Baptista et al., 2015). However, less is known about effects of motor learning and brain stimulation on prefrontal and hippocampal GABA (Hone-Blanchet, Edden, & Fecteau, 2016; Iwabuchi et al., 2017). Considering the limited available literature, our hypotheses with respect to the MRS data are based on M1 studies. Specifically, we hypothesized that facilitatory iTBS and inhibitory cTBS of the DLPFC would result in decreased and increased, respectively, DLPFC and hippocampal GABA levels; and these effects would be more pronounced for sequence learning as compared to the control task. As GABA levels are typically inversely related to BOLD signal (Duncan, Wiebking, & Northoff, 2014), we expected that the intervention-related modulation of activity and connectivity described above will be negatively correlated to the hypothesized changes in DLPFC and hippocampal GABA levels.

## Results

The present research employed an individualized data-driven TBS targeting procedure. To do so, we conducted two experiments on independent participant samples. The goal of Experiment 1 was to identify, using resting-state (RS) fMRI data, a cortical TBS target that was functionally connected to both the hippocampus and striatum. The identified spatial location was then used as the center of a search sphere on individual RS data acquired in Experiment 2 to identify a TBS target significantly and commonly connected to the hippocampus and striatum in each participant. Using this individually-tailored TBS targeting approach, the goal of Experiment 2 was to investigate whether prefrontal stimulation influenced brain responses in cortico-hippocampal and -striatal networks during motor learning.

### Experiment 1: TBS target identification on RS data

To identify a cortical region reachable with TBS and functionally connected to both the striatum and the hippocampus, RS fMRI data of 26 young healthy participants were analyzed. In a first step, seed-based resting-state functional connectivity (RSFC) analyses of the hippocampus and striatum were performed to highlight connectivity patterns of these regions of interest (ROIs) with the rest of the brain. In a second step, a *conjunction analysis* between the hippocampal and striatal RSFC maps was performed in order to identify cortical regions that were significantly and commonly connected to the two ROIs at rest.

As expected, seed-based RSFC analyses using *bilateral hippocampi* as seed regions showed that the hippocampus was significantly (Z ≥ 2.03, *p*_FDR_ < .05) connected, during rest, to a widespread network highly consistent with the default mode network, including the hippocampus, parahippocampus, precuneus, medial prefrontal cortex and posterior cingulate cortex. The connectivity map also highlighted motor-related areas such as the cerebellum, the supplementary motor area and the caudate nucleus (Supplemental Table S1 and Figure 1A left panel). RSFC analyses using *bilateral caudate nuclei* as seed regions showed significant (Z ≥ 1.99, *p*_FDR_ < .05) connectivity during rest with a network that included medial and more lateral frontal areas, but also occipital and temporal regions. Results also showed connectivity at rest with motor-related areas such as the cerebellum, the supplementary motor area and the putamen (Supplemental Table S1 and Figure 1A right panel).

**Figure 1.**
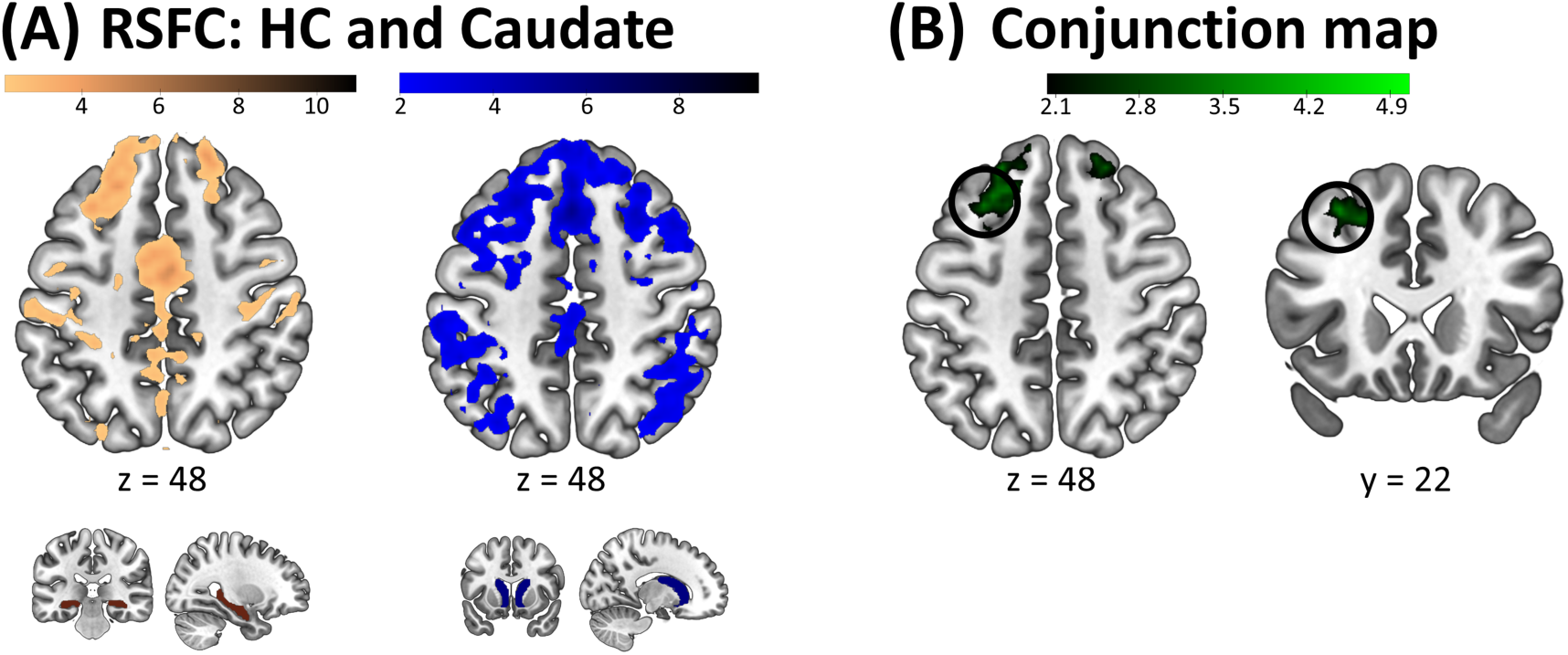
Results of Experiment 1. (A) Resting State Functional Connectivity (RSFC) maps of the hippocampus (HC, left panel) and the caudate nucleus (right panel). The respective seeds are depicted below the connectivity maps. (B) Conjunction map between the HC and Caudate RSFC maps (displayed within a frontal mask). A 15-mm radius sphere (depicted as a black circle) centered around the peak maxima (−30 22 48 mm) was used as search area for individualized targeting in Experiment 2. Connectivity maps and RSFC seeds are displayed on a T1-weighted template image with a threshold of *p_FDR_* < .05 for the connectivity maps. Color bars represent Z values.

**Table 1:**
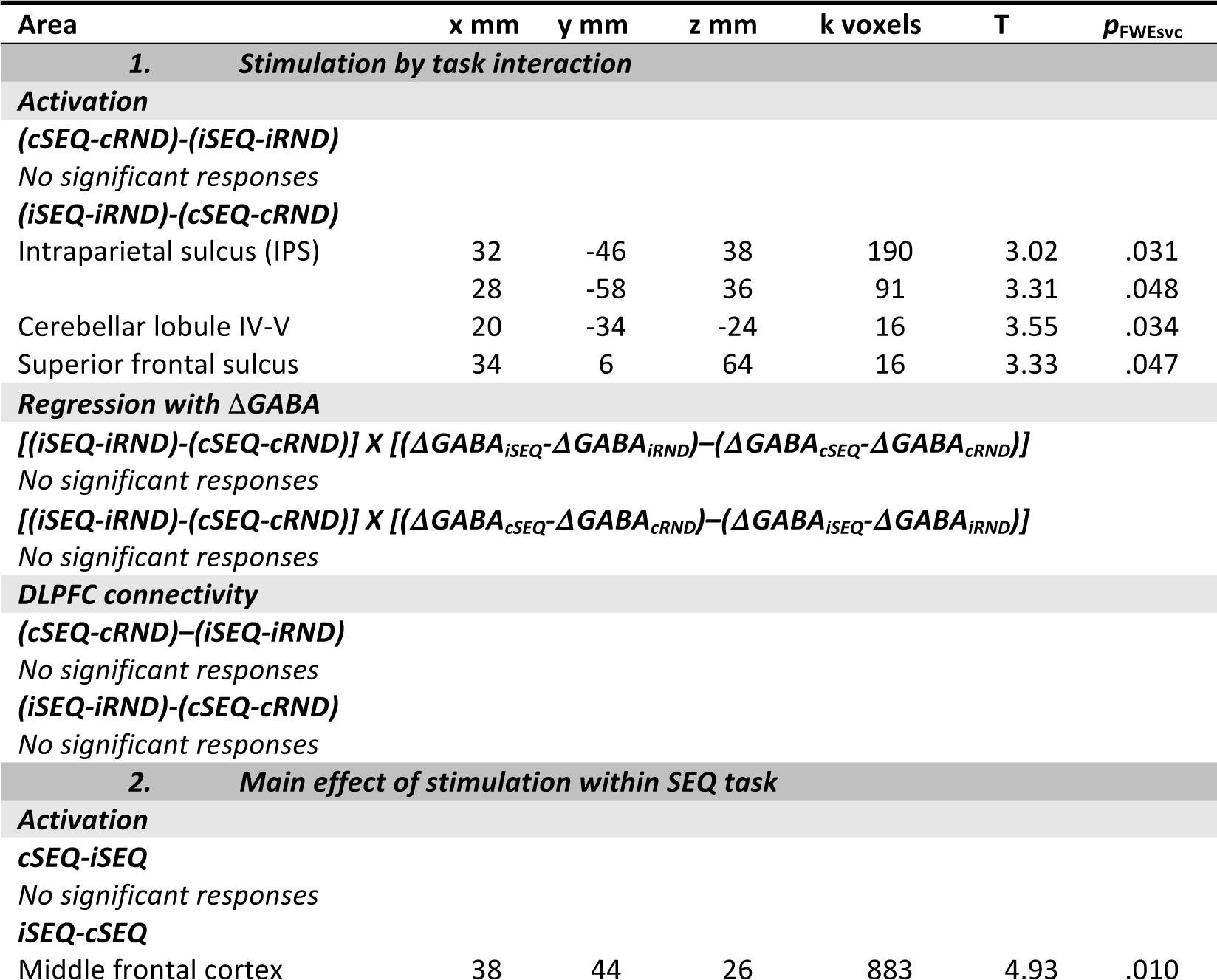
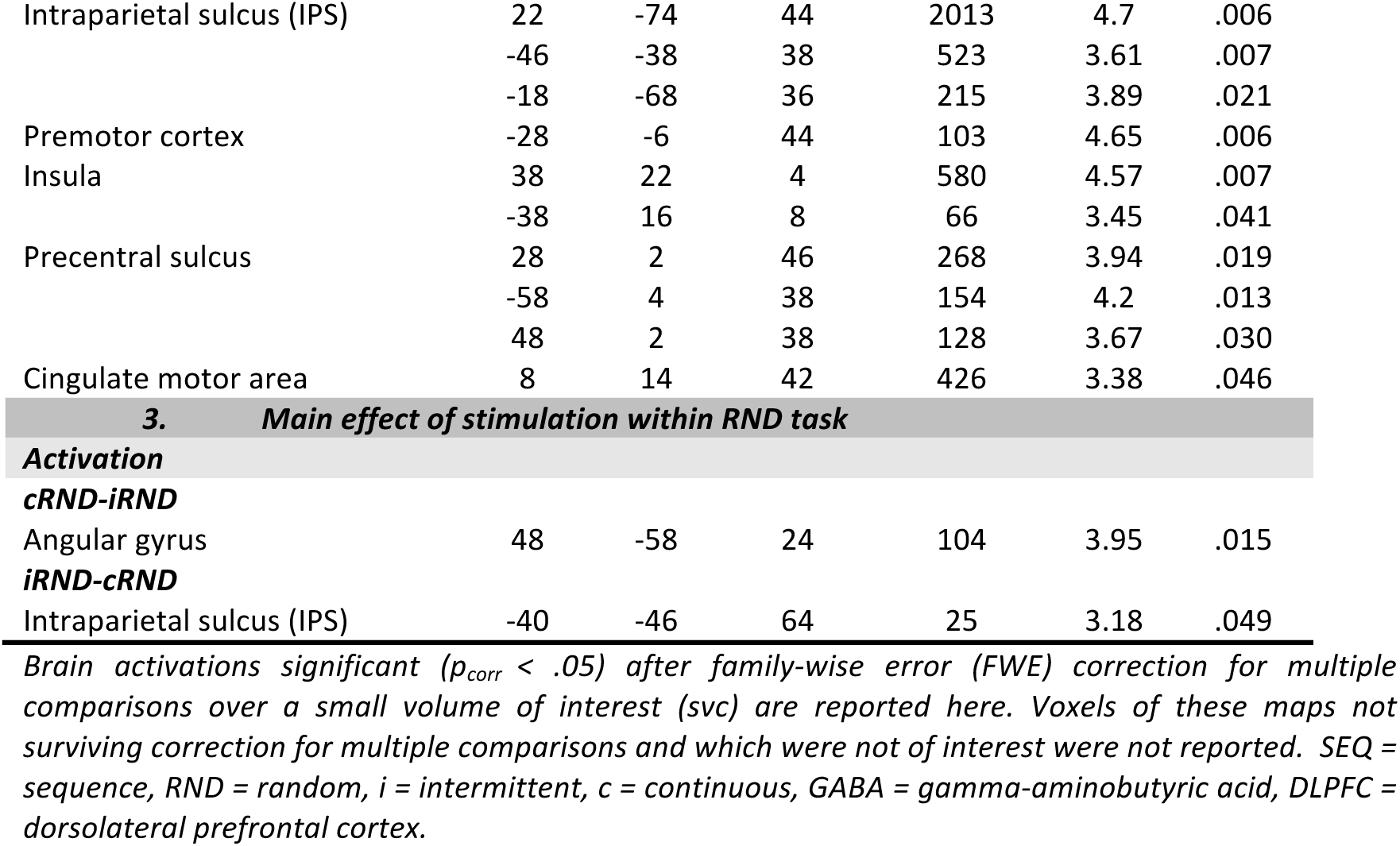
Functional imaging results for the stimulation by task interaction contrasts.

| Area | x mm | y mm | z mm | k voxels | T | $p_{FWESVC}$ |
| --- | --- | --- | --- | --- | --- | --- |
| **1. Stimulation by task interaction** |  |  |  |  |  |  |
| **Activation** |  |  |  |  |  |  |
| **(cSEQ-cRND)-(iSEQ-iRND)** |  |  |  |  |  |  |
| No significant responses |  |  |  |  |  |  |
| **(iSEQ-iRND)-(cSEQ-cRND)** |  |  |  |  |  |  |
| Intraparietal sulcus (IPS) | 32 | -46 | 38 | 190 | 3.02 | .031 |
|  | 28 | -58 | 36 | 91 | 3.31 | .048 |
| Cerebellar lobule IV-V | 20 | -34 | -24 | 16 | 3.55 | .034 |
| Superior frontal sulcus | 34 | 6 | 64 | 16 | 3.33 | .047 |
| **Regression with $\Delta GABA$** |  |  |  |  |  |  |
| **$[(iSEQ-iRND)-(cSEQ-cRND)] \times [(\Delta GABA_{iSEQ}-\Delta GABA_{iRND})-(\Delta GABA_{cSEQ}-\Delta GABA_{cRND})]$** |  |  |  |  |  |  |
| No significant responses |  |  |  |  |  |  |
| **$[(iSEQ-iRND)-(cSEQ-cRND)] \times [(\Delta GABA_{cSEQ}-\Delta GABA_{cRND})-(\Delta GABA_{iSEQ}-\Delta GABA_{iRND})]$** |  |  |  |  |  |  |
| No significant responses |  |  |  |  |  |  |
| **DLPFC connectivity** |  |  |  |  |  |  |
| **(cSEQ-cRND)-(iSEQ-iRND)** |  |  |  |  |  |  |
| No significant responses |  |  |  |  |  |  |
| **(iSEQ-iRND)-(cSEQ-cRND)** |  |  |  |  |  |  |
| No significant responses |  |  |  |  |  |  |
| **2. Main effect of stimulation within SEQ task** |  |  |  |  |  |  |
| **Activation** |  |  |  |  |  |  |
| **cSEQ-iSEQ** |  |  |  |  |  |  |
| No significant responses |  |  |  |  |  |  |
| **iSEQ-cSEQ** |  |  |  |  |  |  |
| Middle frontal cortex | 38 | 44 | 26 | 883 | 4.93 | .010 |
| Intraparietal sulcus (IPS) | 22 | -74 | 44 | 2013 | 4.7 | .006 |
|  | -46 | -38 | 38 | 523 | 3.61 | .007 |
|  | -18 | -68 | 36 | 215 | 3.89 | .021 |
| Premotor cortex | -28 | -6 | 44 | 103 | 4.65 | .006 |
| Insula | 38 | 22 | 4 | 580 | 4.57 | .007 |
|  | -38 | 16 | 8 | 66 | 3.45 | .041 |
| Precentral sulcus | 28 | 2 | 46 | 268 | 3.94 | .019 |
|  | -58 | 4 | 38 | 154 | 4.2 | .013 |
|  | 48 | 2 | 38 | 128 | 3.67 | .030 |
| Cingulate motor area | 8 | 14 | 42 | 426 | 3.38 | .046 |
| cRND-iRND |  |  |  |  |  |  |
| Angular gyrus | 48 | -58 | 24 | 104 | 3.95 | .015 |
| iRND-cRND |  |  |  |  |  |  |
| Intraparietal sulcus (IPS) | -40 | -46 | 64 | 25 | 3.18 | .049 |

Brain activations significant ( $p_{corr} < .05$ ) after family-wise error (FWE) correction for multiple comparisons over a small volume of interest (svc) are reported here. Voxels of these maps not surviving correction for multiple comparisons and which were not of interest were not reported. SEQ = sequence, RND = random, i = intermittent, c = continuous, GABA = gamma-aminobutyric acid, DLPFC = dorsolateral prefrontal cortex.

The *conjunction analysis* performed between the hippocampal and caudate RSFC maps indicated that a network including ventral medial prefrontal, dorsolateral prefrontal, parietal and subcortical regions was significantly *and* commonly connected to both seed regions. Based on evidence reviewed above that (1) the DLPFC plays a pivotal role in the interaction between hippocampal and striatal systems during MSL (Albouy, King, et al., 2013) and that (2) repetitive TMS of the DLPFC can influence brain responses in these deep regions (e.g., Bilek et al., 2013; Ott et al., 2011), we constrained our stimulation target search on the conjunction map to a mask including the middle and superior frontal segments of the AAL atlas (Tzourio-Mazoyer et al., 2002). The resulting statistical map is shown in Figure 1B and the list of identified frontal peaks is presented in Supplemental Table S2. The stimulation target - to be used in Experiment 2 to guide the individualized targeting pipeline - was defined as the peak maxima in the masked conjunction map and was located in the left DLPFC (−30 22 48 mm, encircled in black in Figure 1B).

**Table 2:**
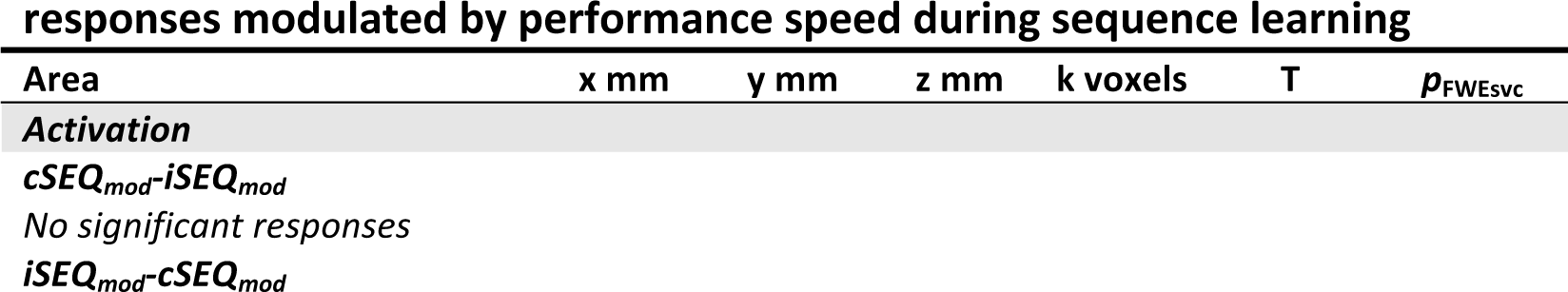
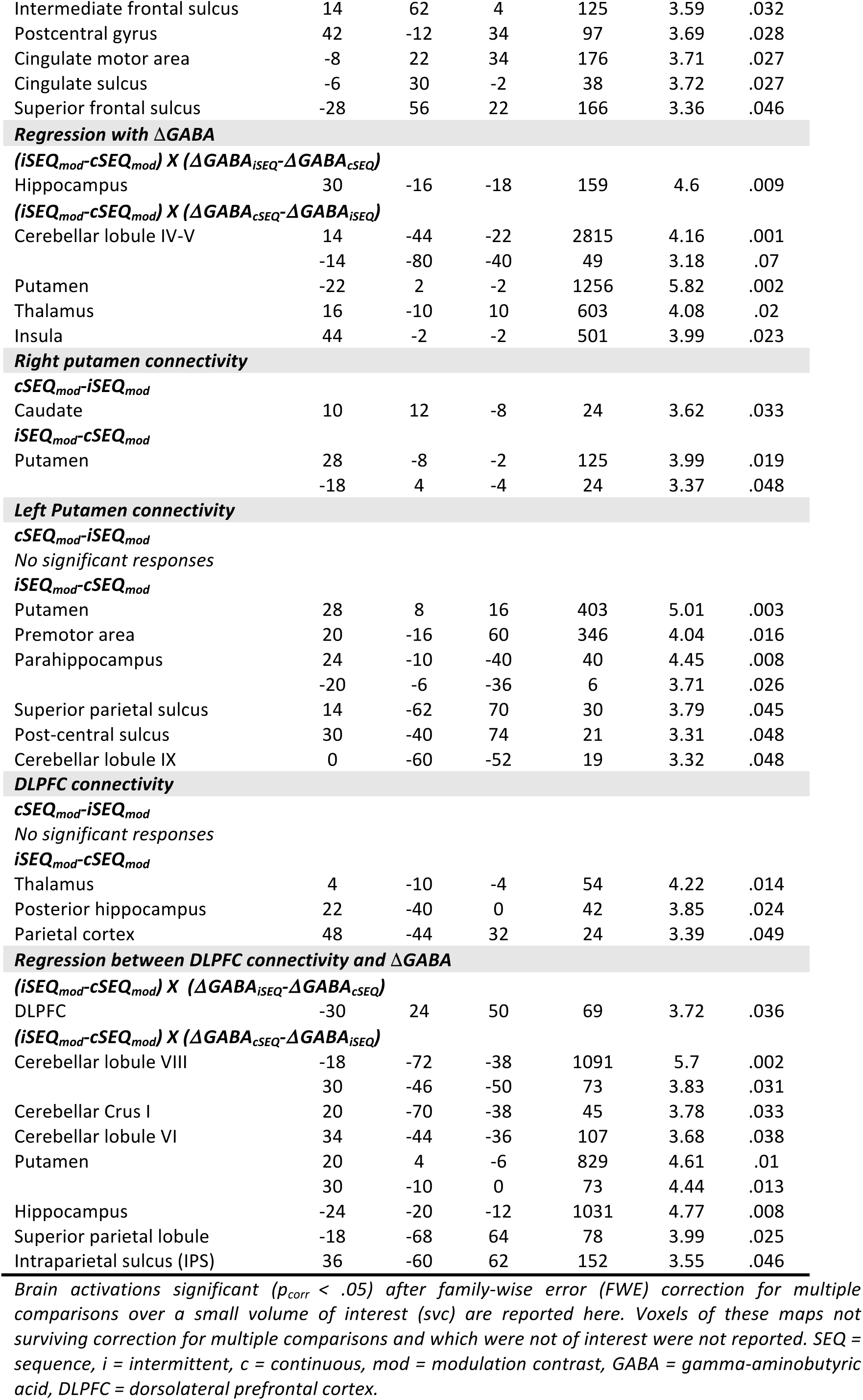
Functional imaging results for the main effect of stimulation on brain responses modulated by performance speed during sequence learning

### Experiment 2: TBS and motor sequence learning

The aim of the second experiment was to investigate the effect of facilitatory iTBS and inhibitory cTBS of the prefrontal target on the neural correlates of motor sequence learning. Individual TBS target definition for the 19 healthy participants included in this experiment was performed using data from a baseline RS fMRI session (see methods). The targeting procedure was similar to that in Experiment 1, but the search was constrained to a 15-mm sphere (Wang et al., 2014) centered on the DLPFC coordinate identified in the first experiment (see Figure 1B). After target definition, participants completed four experimental TBS-MR sessions that were separated by at least six days (i.e. within-subject design; Figure 2). In each session, after receiving T1-neuronavigated facilitatory iTBS or inhibitory cTBS of the individualized DLPFC target, participants performed a sequential (SEQ) or random (RND) version of the serial reaction time task (SRTT) inside the MR scanner. This resulted in four experimental conditions per participant: cTBS/SEQ (cSEQ), cTBS/RND (cRND), iTBS/SEQ (iSEQ) and iTBS/RND (iRND). Magnetic resonance spectroscopy (MRS) data of the DLPFC and the hippocampus were acquired pre-TBS and post-TBS/task.

**Figure 2.**
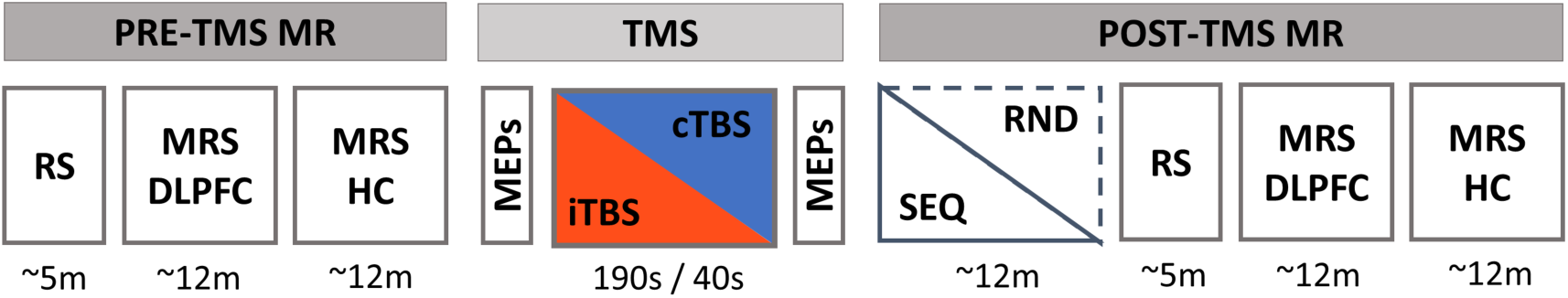
In each experimental session, participants first underwent pre-TMS whole-brain resting-state (RS) fMRI scans and magnetic resonance spectroscopy (MRS) scans of the dorsolateral prefrontal cortex (DLPFC) and the hippocampus (HC) that were followed by T1-neuronavigated intermittent or continuous theta-burst stimulation (iTBS or cTBS) applied to an individually-defined DLPFC target outside the scanner. Motor evoked potentials (MEPs) were measured pre- and post-TBS to probe corticospinal excitability. Immediately following the end of the TMS session, participants were placed in the MR scanner where they were trained on the motor task (sequential [SEQ] or random [RND] versions of the serial reaction time task) while BOLD images were acquired. After task completion, post-TBS/task RS and MRS data of the DLPFC and hippocampus were acquired. The order of the four experimental conditions in this within-subject design [cTBS/SEQ (cSEQ), cTBS/RND (cRND), iTBS/SEQ (iSEQ), iTBS/RND (iRND)] was counterbalanced across participants. Note that the data related to the pre- and post-TBS RS scans are not reported in the present manuscript. TMS: transcranial magnetic stimulation.

### Behavior

The sequential and random SRTT included 16 blocks of practice from which performance speed (i.e., mean reaction time of correct button presses) and accuracy (i.e., percentage of correct button presses) were extracted. Performance speed was faster during the SEQ as compared to the RND task (main effect of task; F_(1,16)_ = 40.435, *p* < .001) and improved over the course of training across task conditions (main effect of block; F_(3.419,54.7)_ = 16.325, *p* < .001). This increase was more pronounced in the SEQ as compared to the RND task (task by block interaction; F_(3.838,61.415)_ = 21.492, *p* < .001; Figure 3, upper panel). No effects of stimulation (F_(1,16)_= 1.639, *p* = .219), stimulation by task (F_(1,16)_ = 2.102, *p* = .166), stimulation by block (F_(7.396,118.341)_ = .446, *p* = .88) or stimulation by task by block (F_(6.155,98.477)_ = .566, *p* = .76) were observed for performance speed.

**Figure 3.**
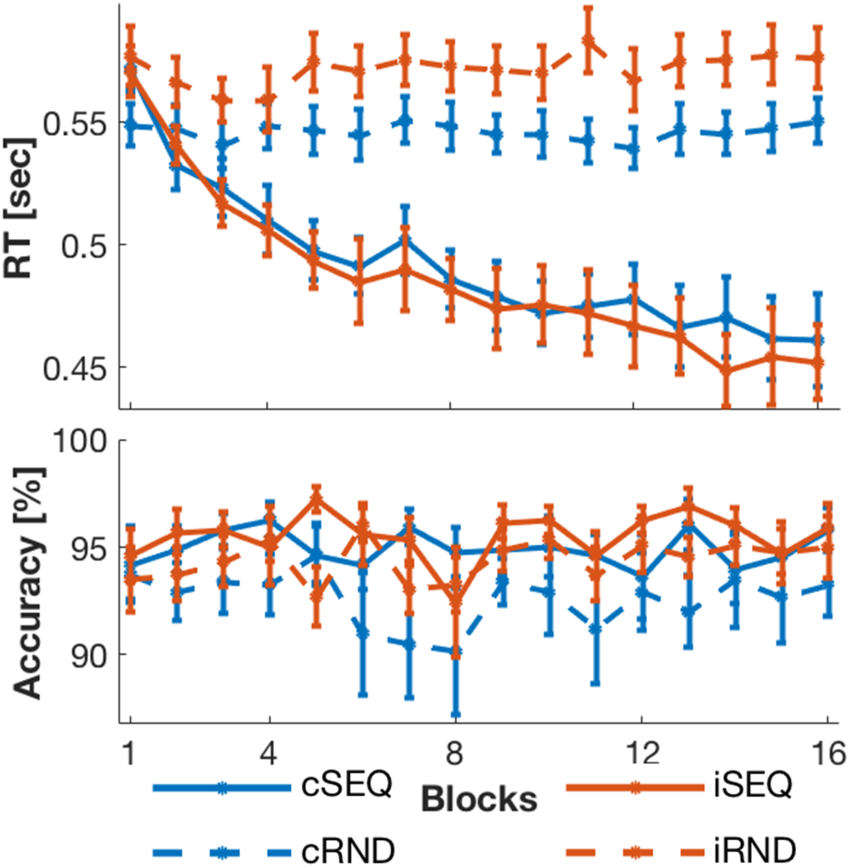
Behavioral results of Experiment 2. Upper panel: Performance speed (reaction time, RT) improved over the course of training in the sequence task (SEQ) conditions and stayed stable in random task (RND) conditions. Lower panel: Performance accuracy remained stable in all conditions with overall higher accuracy in the SEQ than in the RND condition. The stimulation intervention (c: continuous and i: intermittent) did not affect motor performance nor motor learning.

Performance accuracy was higher during SEQ compared to RND practice (main effect of task; F_(1,16)_ = 6.919, *p* = .018; Figure 3, lower panel). No effects of stimulation (F_(1,16)_ = 2.367, *p* = .143), block (F_(4.815,77.033)_ = 1.552, *p* = .186), stimulation by task (F_(1,16)_ = .31, *p* = .585), stimulation by block (F_(5.635,90.163)_ = .662, *p* = .671), task by block (F_(15,240)_ = .643, *p* = .837) or stimulation by task by block (F_(3.476,55.619)_ = .759, *p* = .54) were observed for performance accuracy.

Altogether, the behavioral results demonstrated that participants learned the motor sequence and that the stimulation intervention did not impact motor sequence learning nor overall motor performance.

### MRS of GABA

Fitting of the GABA peak failed in a high proportion of measurements for the hippocampal MRS data, leaving only 10 complete data sets (see methods for further information). As too few measurements remained for appropriate statistical analyses of the hippocampal MRS data, results presented in this paper are limited to the DLPFC voxel (see Figure 4A for a depiction of DLPFC MRS voxel positioning and Supplemental Figure S1 for voxel placements across sessions and participants).

**Figure 4.**
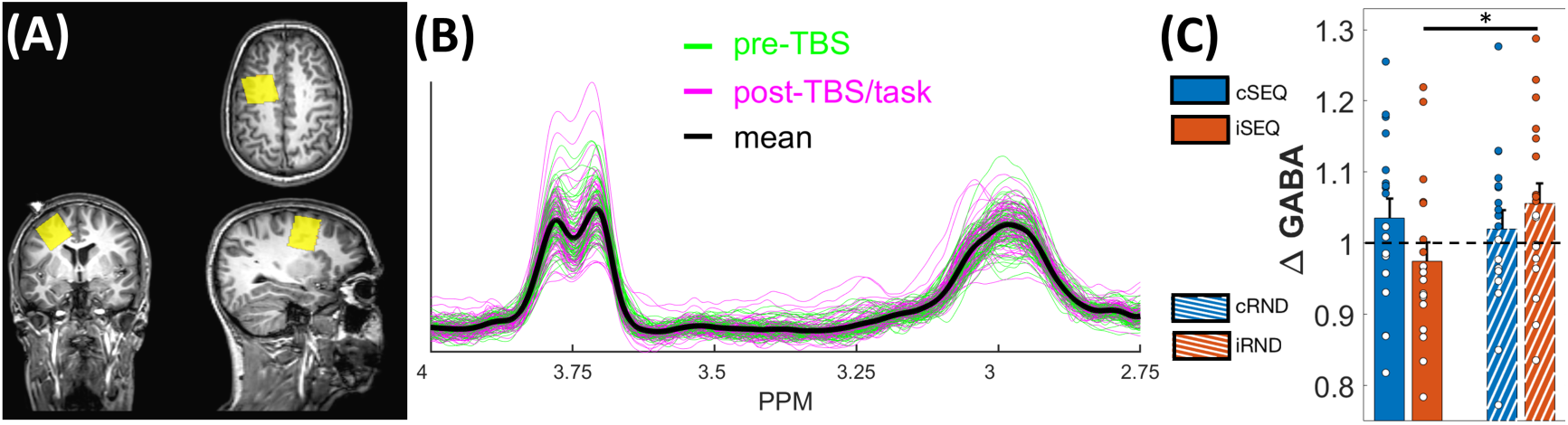
MRS data of the DLPFC voxel. (A) Depiction of DLPFC MRS voxel positioning of a randomly selected participant and time point. The MRS voxel is overlaid on the participant- and time point-specific T1 structural scan. A glycerin maker was placed at the site of stimulation and was used to optimize MRS voxel positioning (marker visible on the coronal view). (B) Spectra of all DLPFC MRS measurements (N = 150), from all participants and time points. GABA+ peak is visible at 3 ppm. Pre-TBS and post-TBS/task time points are depicted in green and magenta, respectively (mean spectrum across all participants and time points depicted in black). (C) ΔGABA in the four experimental conditions. Note that a pre- to post-intervention GABA+ increase and decrease are represented by values above and below 1 (indicated by the black dashed line), respectively. Exploratory analyses indicate that ΔGABA significantly differed between the iSEQ and iRND conditions. Error bars indicate SEM. Circles represent individual data points. Asterisk represents significant paired t-test with *p* < .05. TBS: theta-burst stimulation, i: intermittent, c: continuous, SEQ: sequence, RND: random.

Post-TBS/task GABA+ levels were normalized to pre-TBS GABA+ levels in order to assess intervention-related GABA+ changes (referred to as ΔGABA, see Supplemental Table S3 for raw data and Figure 4B for spectra of all DLPFC MRS measurements). ΔGABA was not significantly influenced by the task (F_(1,16)_ = 2.181, *p* = .159), stimulation (F_(1,16)_ = .025, *p* = .876) or by an interaction between task and stimulation (F_(1,16)_ = 2.975, *p* = .104). Exploratory paired t-tests indicated that GABA+ levels were significantly reduced after sequence learning as compared to random practice under the influence of iTBS (iSEQ vs. iRND; t_(1,17)_ = −2.508, *p* = .023; Figure 4C). None of the other paired comparisons were significant (all *ps* > .05).

**Table 3:** Participant characteristics of Experiment 2

|  |  |
| --- | --- |
| *N* | 19 (12 females) |
| *Age (years)* | 22.42 ($\pm 2.36$) |
| *Beck Anxiety Inventory* | 1.68 ($\pm 2.43$) |
| *Beck Depression Inventory* | 3.84 ($\pm 3.95$) |
| *Edinburgh Handedness* | 85.26 ($\pm 14.57$) |
| *Pittsburgh Sleep Quality Index* | 3.52 ($\pm 1.54$) |
| *St. Mary Quality of sleep* | 4.04 ($\pm 0.8$) |
| *St. Mary Quantity of sleep* | 8h ($\pm 1.01$) |
| *Stanford Sleepiness Scale* | 2.11 ($\pm 0.76$) |

### Functional brain imaging data

We investigated the effects of stimulation and task conditions on the amplitude and the dynamics of task-related activity and connectivity. Connectivity was assessed with psychophysiological interaction (PPI) analyses using, as seed regions, the DLPFC (TBS target) and any striatal and hippocampal region highlighted in task-related activity contrasts (see methods). Additionally, we performed regression analyses between ΔGABA and the above-mentioned activity and DLPFC connectivity maps to assess the relationships between changes in prefrontal GABA pre- to post-intervention and BOLD responses during task performance.

### Stimulation by task interaction Activity

We first analyzed the stimulation by task interaction contrast averaged across the 16 blocks of practice (Table 1.1). The intraparietal sulcus (IPS), the cerebellar lobule and the frontal cortex were more activated during sequence learning as compared to random practice after iTBS than cTBS (*(iSEQ-iRND)-(cSEQ-cRND)*, Table 1.1). These effects were mainly driven by larger differences between stimulation conditions in the SEQ as compared to the RND task condition (Figure 5A). Indeed, follow-up comparisons between stimulation conditions within the SEQ task indicated that a large network including frontal areas, IPS and insula showed higher activity after iTBS compared to cTBS (*iSEQ-cSEQ*, Table 1.2; Figure 5B). There was only a limited number of areas showing a significant stimulation effect within the RND task (*cRND vs. iRND*, Table 1.3). These results collectively indicate that the two stimulation conditions had a differential impact on activity in fronto-parietal-cerebellar areas specifically during sequence learning.

**Figure 5.**
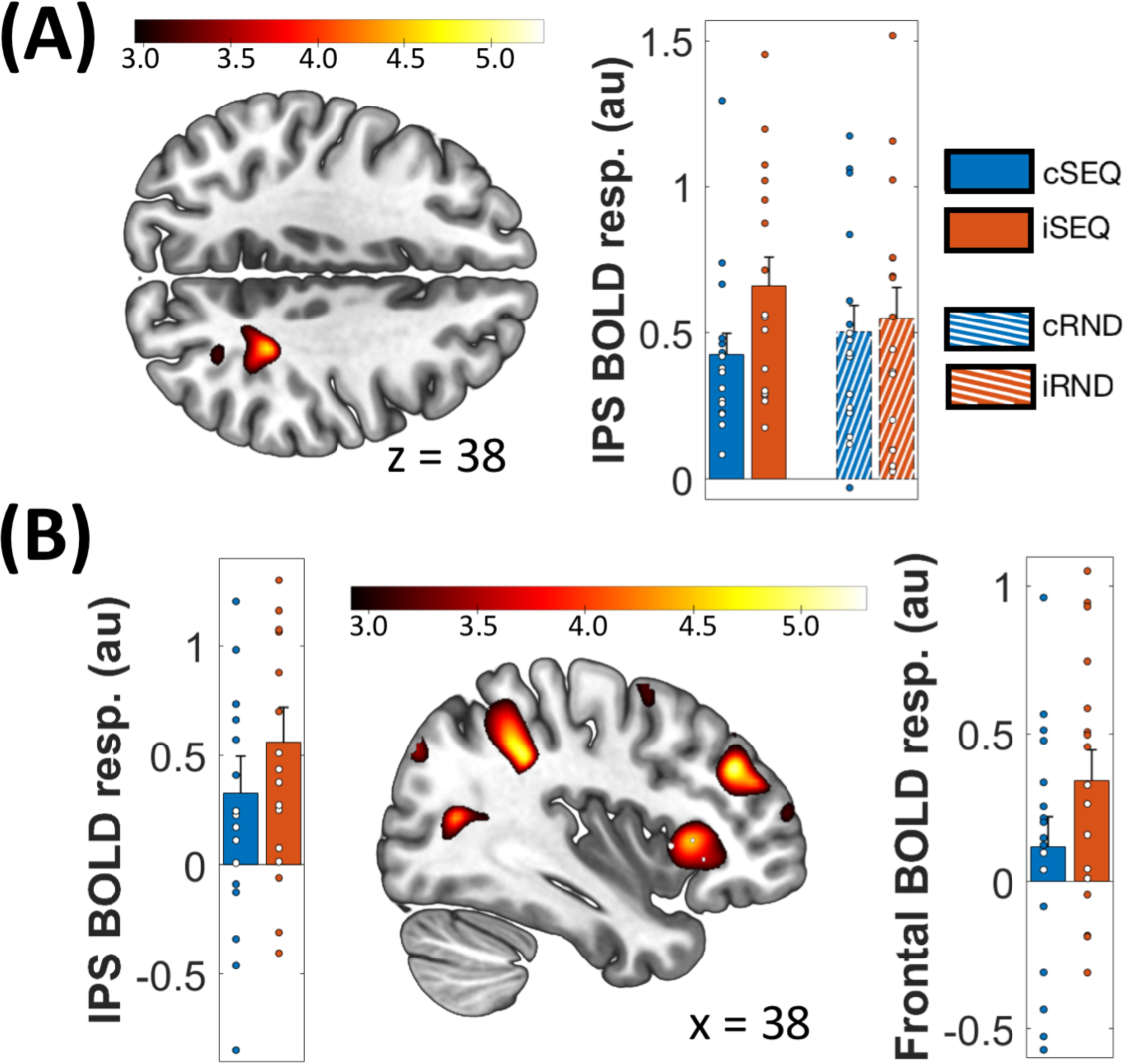
(A) Stimulation by task interaction in the right intraparietal sulcus (IPS, 32 −46 38 mm). IPS was more activated during SEQ as compared to RND after iTBS than cTBS. (B) Stimulation effect within SEQ task in the IPS (22 −74 44 mm), insula (38 22 4 mm) and frontal cortex (38 44 26 mm) was driven by higher activity after iTBS than cTBS. Activations maps are displayed on a T1-weighted template image with a threshold of *p* < .005 uncorrected. Color bars represent T values. Error bars indicate SEM. Circles represent individual data points. au: arbitrary units, resp.: response, TBS: theta-burst stimulation, i: intermittent, c: continuous, SEQ: sequence, RND: random.

Regression analyses between the interaction contrast reported above and changes in GABA+ across the two time points (ΔGABA) in the four experimental conditions did not reveal any significant results.

#### Connectivity

As the stimulation by task interactions reported above did not reveal any significant clusters in the striatum and hippocampus, connectivity analyses with seeds from these two regions of interest were not performed. Analyses assessing whether connectivity of the DLPFC stimulation target was influenced by an interaction between stimulation and task conditions did not show any significant results.

#### Learning-related modulation of brain responses

We used parametric modulation analyses (see methods) to test whether brain activity changed as a function of learning, i.e. the block-to-block performance improvements, in the SEQ conditions. This allowed us to examine whether the different stimulation conditions influenced the learning-related *dynamics* of brain responses.

#### Activity

Consistent with previous research (Albouy et al., 2012), activity in bilateral putamen, as well as in the cerebellum, increased as a function of learning regardless of the type of stimulation (i.e., *iSEQ_mod_+cSEQ_mod_*, Supplemental Table S4).

Between-stimulation-condition contrasts showed that a set of brain regions, including superior frontal areas, central sulcus and cingulum, were differently modulated by learning depending on the stimulation condition (*iSEQ_mod_-cSEQ_mod_*, Table 2). These effects were driven by a progressive learning-related increase in activity in the aforementioned regions in cSEQ over the course of training (Supplemental Figure S2).

We then conducted regression analyses assessing whether between-condition differences in dynamical brain activity during training were related to differences in DLPFC ΔGABA between conditions. Results show that a between-condition difference (*iSEQ_mod_-cSEQ_mod_*) in dynamical activity in the hippocampus was related to the difference in DLPFC ΔGABA between stimulation conditions (*ΔGABA_iSEQ_-ΔGABA_cSEQ_*, Table 2; Figure 6A). Interestingly, between-condition differences in dynamical activity in putamen and cerebellar activity were also related to the difference in DLPFC ΔGABA between stimulation conditions, but in the opposite direction (*ΔGABA_cSEQ_-ΔGABA_iSEQ_*) as compared to the hippocampus. It is worth noting that, given our within-subject design and the corresponding appropriate statistical models, it is not possible to provide a more directional interpretation of these patterns of results (see discussion for further details).

**Figure 6.**
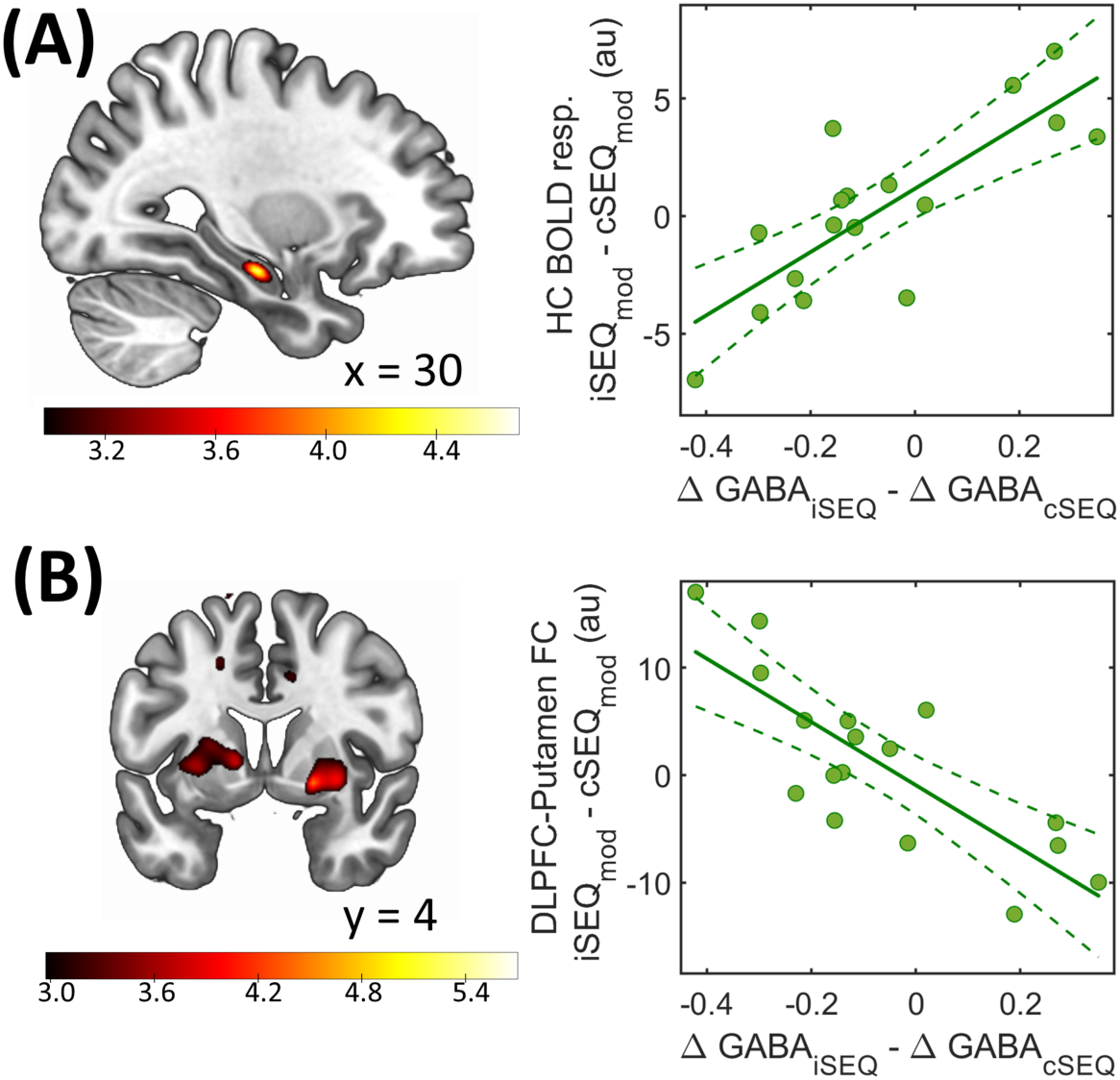
Regressions with DLPFC ΔGABA. (A) Hippocampal (HC) dynamical activity during learning (30 −16 −18 mm, left panel) was differently related to DLPFC ΔGABA between conditions. (B) Learning-related changes in DLPFC-putamen functional connectivity (FC) patterns (20 4 −6 mm, left panel) were differently related to DLPFC ΔGABA between conditions. Regression maps are displayed on a T1-weighted template image with a threshold of *p* < .005 uncorrected. Color bars represent T values. Circles represent individual data, solid lines represent linear regression fits, dashed lines depict 95% prediction intervals of the linear function. au: arbitrary units, resp.: response, i: intermittent, c: continuous, SEQ: sequence, mod: modulation contrast, GABA = gamma-aminobutyric acid.

In sum, these results indicate that the DLPFC stimulations differently influenced (1) the dynamical learning-related patterns of activity in the central sulcus as well as frontal and cingulate areas; and, (2) the relationship between changes in DLPFC GABA levels and learning-related changes in activity patterns in the hippocampus, striatum and cerebellum.

#### Connectivity

Connectivity analyses were performed using, as seed regions, the putamen clusters described above that exhibited increases in activity as a function of learning across the two stimulation conditions (Supplemental Table S4). Functional connectivity between these bilateral putamen seeds and sensorimotor parts of the putamen increased as a function of learning more in the iSEQ as compared to the cSEQ condition (*iSEQ_mod_-cSEQ_mod_*, Table 2; Figure 7A upper panel). In contrast, the right putamen showed a greater learning-related increase in connectivity with the caudate nucleus, a more associative territory of the striatum, in the cSEQ as compared to the iSEQ condition (*cSEQ_mod_-iSEQ_mod_*, Table 2; Figure 7A lower panel).

**Figure 7.**
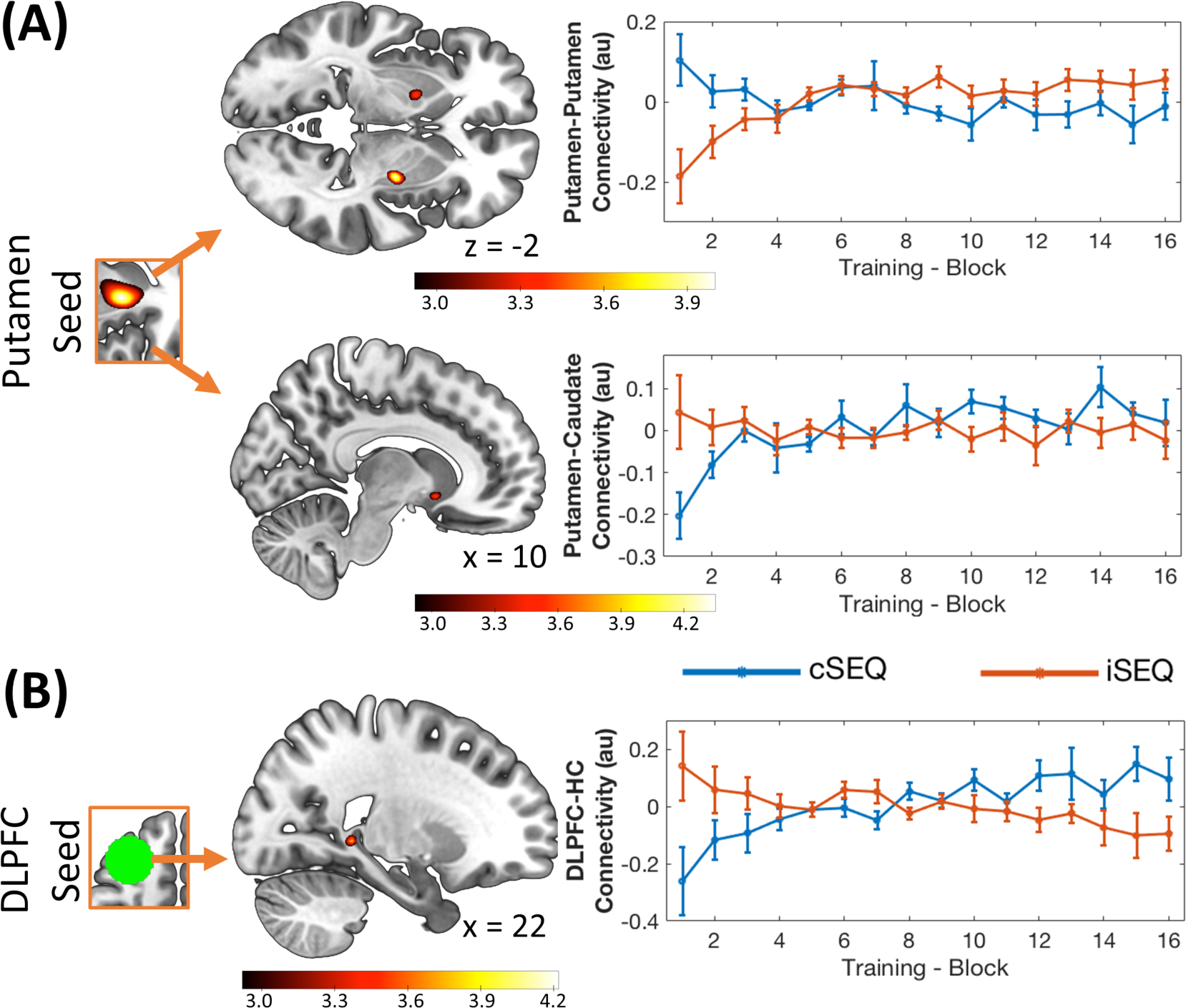
Stimulation effect on sequence (SEQ) task-related connectivity. (A) Functional connectivity (FC) between the right putamen and the sensorimotor putamen (28 −8 −2 mm, upper panel) increased more as a function of learning after iTBS compared to cTBS. FC with the caudate nucleus (10 12 −8 mm, lower panel) showed the opposite pattern. (B) FC of the DLPFC TBS target with the hippocampus (HC, 22 −40 0 mm) increased more as a function of learning in the cTBS as compared to the iTBS condition. Connectivity maps are displayed on a T1-weighted template image with a threshold of *p* < .005 uncorrected. Color bars represent T values. Error bars indicate SEM. au: arbitrary units, TBS: theta-burst stimulation, i: intermittent, c: continuous.

Functional connectivity analyses using the DLPFC TBS target as a seed region indicate that the dynamical connectivity patterns between the DLPFC and a set of areas including the thalamus, hippocampus and the fusiform area were different between stimulation conditions. These differences in fronto-hippocampal connectivity were explained by antagonistic dynamical patterns between conditions; specifically, connectivity decreased and increased as a function of learning in the iTBS and cTBS conditions, respectively (*iSEQ_mod_-cSEQ_mod_* contrast, Table 2; Figure 7B).

Regression analyses linking between-condition differences in DLPFC connectivity (*iSEQ_mod_-cSEQ_mod_*) to ΔGABA (*ΔGABA_iSEQ_ vs. ΔGABA_cSEQ_*) showed that the dynamical connectivity patterns between the DLPFC and a set of brain regions including the prefrontal cortex, the putamen, hippocampus and the cerebellum were differently related to the DLPFC ΔGABA between stimulation conditions (Table 2, Figure 6B). Similar to the results presented on activation contrasts, these correlations could be explained by various individual patterns of connectivity modulation and ΔGABA (see discussion).

Altogether, our results indicate that iTBS, as compared to cTBS, applied to the DLPFC before motor sequence learning promoted learning-related increases in connectivity in sensorimotor-striatal networks. In contrast, cTBS of the DLPFC resulted in progressive connectivity increases in fronto-hippocampal and associative-striatal networks. Additionally, our findings show that the stimulation conditions differently altered the relationship between the learning-related changes in DLPFC-striatum-hippocampus connectivity and DLPFC ΔGABA.

## Discussion

In this proof-of-concept study, we investigated whether non-invasive brain stimulation of the prefrontal cortex can alter neural responses in hippocampal and striatal networks during motor sequence learning. Specifically, we employed a data-driven approach to first identify a prefrontal target region that was functionally connected to the striatum and the hippocampus at rest. This target was subsequently used to guide an individualized TBS targeting procedure in a separate TBS-MRI experiment. The results of this second experiment showed that facilitatory iTBS, as compared to inhibitory cTBS, applied before sequence learning, as compared to random practice, induced greater task-related activity in fronto-parieto-cerebellar regions. Interestingly, the different stimulation conditions also altered the dynamical connectivity patterns in fronto-hippocampal and striatal networks during learning. While facilitatory iTBS promoted learning-related increases in connectivity in sensorimotor-striatal networks, inhibitory cTBS resulted in progressive connectivity increases in fronto-hippocampal and associative-striatal networks. Finally, facilitatory iTBS and inhibitory cTBS differently influenced the relationship between changes in DLPFC GABA+ levels and both dynamical activity and connectivity patterns of the hippocampus and striatum during motor sequence learning. This research is, to the best of our knowledge, the first to demonstrate that brain stimulation can influence motor learning-related responses in the striatum and the hippocampus.

### DLPFC stimulation altered activity in fronto-parietal-cerebellar regions during motor sequence learning

Previous research investigating the effect of DLPFC stimulation on brain activity *at rest* or during *task-practice* has reported significant effects on the brain responses in the stimulated area itself as well as in widespread networks including various cortical areas (motor, frontal, parietal, cingulate, temporal, insula), the cerebellum and deep regions including the striatum and the hippocampus (Esslinger et al., 2014; Gratton, Lee, Nomura, & D’Esposito, 2014; Hanlon et al., 2013, 2016; Rounis et al., 2006; Shang et al., 2019; Tang et al., 2019; van der Werf et al., 2010; Xue et al., 2017). Our results indicate that facilitatory iTBS, as compared to inhibitory cTBS, applied prior to motor sequence learning, as compared to random, induced greater task-related activity in the cerebellum and the parietal cortex as well as in frontal areas non-overlapping with the TBS target. Importantly, these brain regions are known to be critical for motor sequence learning. Specifically, cerebellar, parietal as well as frontal regions have been described to support the early motor learning process (Albouy, King, et al., 2013; Doyon et al., 2009, 2018; Hikosaka et al., 2002). Specifically, parieto-frontal loops, with associative territories of the cerebellum and the basal ganglia, are thought to process the spatial (effector-independent) representation of the motor sequence under high control and attentional processes during initial learning. Our results therefore suggest that the two prefrontal stimulation conditions differently altered the control processes supporting these early representations of the motor sequence.

Additionally, we observed that facilitatory iTBS, as compared to inhibitory cTBS, elicited greater activity in the insula, albeit this effect was not sequence learning specific (i.e., not significantly larger in the sequence as compared to the random condition). These findings are in line with previous reports of DLPFC stimulation-induced effects on insular activity and connectivity (Gratton, Lee, Nomura, & D’Esposito, 2013; Hanlon et al., 2016; Iwabuchi et al., 2017; Rounis et al., 2006). Based on evidence that the insula is part of the fronto-parietal attentional network (Fox et al., 2005; Toro, Fox, & Paus, 2008), we propose that the influence of stimulation on activity in fronto-insular-parietal areas might have influenced spatial attention and working memory processes that are typically observed early during task practice (Hikosaka et al., 2002). As these effects were not sequence-specific, attentional processes appeared to have been altered in the random version of the task as well.

### DLPFC stimulation influenced connectivity in fronto-hippocampal and striatal networks during motor sequence learning

Previous research has shown that repetitive TMS of the DLPFC can alter functional connectivity patterns of the target region with other cortical areas (e.g., the insula, cingulate, parietal and frontal cortices), as well as with deep regions including the striatum and the hippocampus, during *resting-state* (Alkhasli et al., 2019; Esslinger et al., 2014; Gratton et al., 2013; Iwabuchi et al., 2017; Mastropasqua et al., 2014; Shang et al., 2019) and during working and episodic memory *tasks* (Bilek et al., 2013; Davis, Luber, Murphy, Lisanby, & Cabeza, 2017; Esslinger et al., 2014). In line with this earlier work, our connectivity analyses revealed that DLPFC stimulation before learning altered connectivity in fronto-hippocampal and striatal networks during motor sequence learning.

Connectivity analyses using the stimulated DLPFC as a seed region indicated that after inhibitory cTBS and facilitatory iTBS, fronto-hippocampal connectivity increased and decreased, respectively, as a function of sequence learning. Interestingly, both activity and connectivity in hippocampo-frontal networks are usually described to decrease as a function of learning under normal (i.e., non-stimulated) conditions (Albouy, King, et al., 2013; Albouy et al., 2008, 2012; Doyon et al., 2018). Our data therefore suggest that inhibitory cTBS disrupted the usually observed pattern of hippocampo-frontal responses during learning. Based on models proposing that the hippocampus, together with the fronto-parietal networks, supports early representations of motor sequences, our connectivity results suggest that inhibitory cTBS might have altered the early control processes discussed above. Importantly, we showed in previous studies that hippocampal activity and connectivity patterns during initial motor sequence learning are critically linked to subsequent consolidation processes (Albouy, King, et al., 2013; Albouy et al., 2008). It is therefore tempting to speculate that the stimulation-induced modulation of hippocampo-fronto-parietal responses might influence subsequent motor memory retention. While this remains hypothetical, it is indeed in line with earlier behavioral work showing that DLPFC stimulation can influence motor memory consolidation (Galea, Albert, Ditye, & Miall, 2010; Tunovic, Press, & Robertson, 2014).

Striatal connectivity analyses indicate that facilitatory iTBS and inhibitory cTBS of the DLPFC promoted a progressive increase in connectivity within sensorimotor- and associative-striatal networks, respectively. Interestingly, this suggests that the different types of stimulation affected functional responses in different striatal networks during motor learning. Previous research has extensively described dynamical activity and connectivity patterns in striatal circuits during sequence learning (Albouy, King, et al., 2013; Doyon et al., 2009; Hikosaka et al., 2002). Task practice is usually paralleled by a gradual shift in activity from associative territories of the striatum (including the caudate nucleus) which support slow and variable performance early during learning (Albouy et al., 2012; Lehéricy et al., 2005), to sensorimotor areas of the putamen when performance plateaus and automatization is reached (Lehéricy et al., 2005). Interestingly, the present results suggest that facilitatory iTBS to the DLPFC further promoted the practice-related shift to sensorimotor striatal functioning. In contrast, inhibitory cTBS altered the usually observed *decrease* in associative striatum involvement and induced learning-related *increases* in connectivity between the associative striatum (caudate nucleus) and the putamen. Together with the observation of inhibitory-stimulation-induced increases in fronto-hippocampal connectivity over the course of learning, the present results indicate that inhibitory prefrontal cTBS promoted the progressive engagement of networks involved in early learning and control processes. It is worth noting that we did not observe any stimulation-induced changes in hippocampo-striatal functional connectivity, such as a decrease in competition between the two networks, as recently proposed by Freedberg et al. (2020).

### DLPFC stimulation altered the relationship between DLPFC GABA+ levels and functional responses in the hippocampus and striatum

MRS data indicated that neither stimulation nor task conditions impacted DLPFC GABA+ levels. However, exploratory analyses within the facilitatory iTBS condition showed larger GABA+ decreases after sequential as compared to random task practice. These results suggest that, under the effect of excitatory stimulation, motor sequence learning, in comparison to random motor execution, resulted in a decrease in DLPFC inhibitory tone. Interestingly, these results are in line with evidence of learning-induced decreases in M1 GABA levels during (Kolasinski et al., 2018), immediately after (Floyer-Lea et al., 2006) but also after 6 weeks of motor sequence learning (Sampaio-Baptista et al., 2015). Importantly and consistent with our results, these M1 GABA changes have been described to be learning-specific, as no such effects were observed after random task practice (Floyer-Lea et al., 2006; Kolasinski et al., 2018). We argue that, similar to M1, the decrease in DLPFC GABA+ levels might reflect disinhibition processes that promote successful learning (Kolasinski et al., 2018; Stagg et al., 2011).

It is worth noting that the learning-specific GABA+ effect in the DLPFC was observed under the effect of facilitatory stimulation, which might suggest that stimulation potentiated the neural plasticity processes mentioned above. Although this interpretation is speculative given the absence of a stimulation by task interaction, it is in line with previous studies describing increases and decreases of M1 GABA levels after inhibitory [e.g., cTBS or cathodal transcranial direct current stimulation (tDCS); (Bachtiar et al., 2018; Stagg, Wylezinska, et al., 2009)] and facilitatory [e.g. anodal tDCS; (Bachtiar et al., 2018, 2015; Stagg, Best, et al., 2009)] stimulation of M1, respectively.

Interestingly, our BOLD / GABA regression analyses showed that the type of stimulation applied before motor sequence learning affected the relationship between DLPFC GABA changes and functional responses in the striatum and the hippocampus. Specifically, the stimulation conditions differently altered the relationship between changes in DLPFC GABA+ levels and learning-related changes in (1) activity patterns in the hippocampus, striatum and cerebellum and (2) in DLPFC-striatum-hippocampus connectivity. These results provide direct support for a central role of the DLPFC in orchestrating the interaction between hippocampal and striatal systems during motor sequence learning (Albouy, King, et al., 2013). Importantly, the present data offer the first evidence that dynamical activity patterns of the hippocampus and striatum as well as fronto-hippocampo-striatal connectivity are related to the changes in inhibitory tone of the DLPFC. Our results also highlight the critical concept that the relationship between DLPFC GABA changes and functional responses in the hippocampus and the striatum can be altered by DLPFC stimulation. However, it is worth noting that, given the nature of the metrics used in the regression models (see ‘Methodological considerations’ section for details), it is not possible to provide a more directional interpretation of these patterns of results, as the BOLD / GABA regression results could reflect various individual patterns. Nonetheless, the findings indicate that, at the group level, differences in learning-related modulations of brain responses in the hippocampus and the striatum were differentially related to changes in DLPFC GABA+ between stimulation conditions.

### DLPFC stimulation did not affect motor performance

The stimulation-induced modulations of activity and connectivity patterns described above did not influence motor performance. Interestingly, the lack of a behavioral difference demonstrates that effects observed at the brain level do not consistently influence motor behavior. Related, the absence of a behavioral effect allows us to attribute the observed differences in functional activity and connectivity, and their relationships to changes in prefrontal GABA levels, directly to the stimulation interventions. In contrast, if TBS impacted behavior, it would not have been possible to rule out the possibility that the stimulation interventions altered behavior, which, in turn, influenced functional activity and connectivity patterns.

It is worth explicitly stating, however, that this null behavioral result may be considered surprising based on the available literature. Previous behavioral work showed that disruptive DLPFC stimulation applied *before* or *during* motor sequence learning can effectively impair motor performance and learning processes (Burke & Coats, 2016; Dayan, Herszage, Laor-Maayany, Sharon, & Censor, 2018; Pascual-Leone, Wassermann, Grafman, & Hallett, 1996; Robertson, Tormos, Maeda, & Pascual-Leone, 2001). The discrepancy between these findings and our current results could be explained by several factors, including differences in stimulation procedure (e.g., TBS vs. 1 Hz, 5 Hz repetitive TMS or single pulse TMS), task complexity (bimanual vs. unimanual tasks), awareness of the sequential material to learn (explicit vs. implicit) and whether reward was provided or not during learning. It is also possible that the absence of behavioral effects in the current study could be the result of compensatory brain responses taking place during task practice after inhibitory prefrontal stimulation. Indeed, based on the neuroimaging results described above, one could have expected that, in the inhibitory stimulation condition, the prolonged engagement of associative striatal and fronto-hippocampal networks - usually observed early during learning when performance is poor – would result in slower performance. As no differences in motor behavior were observed between stimulation conditions, we propose that the sustained engagement of associative striatum-hippocampo-frontal areas during learning under inhibitory cTBS might represent a compensatory mechanism allowing performance to be maintained over the course of practice. The continued engagement of these regions may have counteracted the disruptive effect of stimulation on frontal control processes early during learning and thus may have contributed to improvements in performance during task practice despite a progressive decrease in connectivity within sensorimotor-striatal territories.

### Methodological considerations

It is worth acknowledging that the present study included two active stimulation conditions (i.e., iTBS and cTBS) rather than a sham stimulation. We made the methodological choice to prioritize the inclusion of a control (random) task condition in order to investigate sequence learning-specific effects. The control condition did afford us with the opportunity to test whether the effect of stimulation on brain function depends on the “state” under which stimulation was active (i.e., learning vs. control). A discussion of our results in the context of a no stimulation condition was therefore limited to qualitative comparisons with the available literature.

With respect to the MRS data, it is worth mentioning that GABA+ levels cannot be clearly assigned to one of the various pools of GABA found in the brain [see (Stagg, 2014)]. There is indeed an ongoing scientific discussion in the field, which raises questions about the interpretations of MRS GABA data. Furthermore, due to issues with data quality, we were not able to investigate the effects of our intervention on GABA+ levels in the hippocampus. Additionally, we did not include measurements of striatal GABA due to time constraints imposed by the experimental design. Given the critical roles of these structures in motor sequence learning, it would be of interest for future research to examine learning- and stimulation-induced effects on striatal and hippocampal GABA. Last, and perhaps most importantly, given our within-subject design and the corresponding statistical models necessary to investigate the relationship between BOLD and GABA+ data, it is not possible to provide a directional interpretation of the regression results. Specifically, a significant effect in such an analysis represents a between stimulation condition difference in the relationships between: a) learning-dependent modulations in brain activity/connectivity (referred to as differential modulation betas; depicted on the y-axes on Figure 6); and, b) ΔGABA across the stimulation/task interval (i.e., differential ΔGABA; x-axes on Figure 6). As the beta estimates representing the modulation in brain activity/connectivity as well as ΔGABA are both bi-directional (i.e., values represent an increase or decrease in activity/connectivity with learning or an increase or decrease in GABA+ after the intervention), the difference between stimulation conditions computed on these parameters could then reflect various individual patterns. For example, a large differential modulation beta could be attributed to a steeper decrease in activity in iSEQ than cSEQ or to no modulation in iSEQ and an increase in activity in cSEQ. Similarly, a small differential ΔGABA value could be due to a larger GABA+ decrease in the iSEQ than in the cSEQ condition or to no change in iSEQ and an increase in the cSEQ condition. A deeper inspection of these various possibilities revealed no single pattern that could adequately summarize the reported effects.

## Conclusions

In the present proof-of-concept study that employed a multimodal neuroimaging approach, we demonstrated, for the first time, that DLPFC stimulation influenced *connectivity* patterns within hippocampo-frontal and striatal networks during motor sequence learning. Our data also showed that non-invasive brain stimulation altered the relationship between the levels of inhibition, as assessed with MRS of GABA, in the stimulated area and learning-related changes in both *activity* and *connectivity* in fronto-striato-hippocampal networks. This provides the first experimental evidence that prefrontal brain stimulation can alter functional responses in the striatum and hippocampus during motor learning.

## Methods

The present research included two experiments performed on independent samples. The goal of Experiment 1 was to identify, using resting-state (RS) fMRI data, a TBS target within the prefrontal cortex that was significantly and commonly functionally connected to the hippocampus and striatum. The goal of Experiment 2 was to investigate the effect of facilitatory iTBS and inhibitory cTBS of the prefrontal target, defined individually around the group coordinate identified in Experiment 1, on the behavioral and neural correlates of motor sequence learning.

### Ethics statement

The two experiments presented in this paper were approved by the local Ethics Committee (UZ / KU Leuven). All participants gave their written informed consent before taking part in the study and were compensated for their participation. Procedures were executed in conformity with the approved guidelines.

### Experiment 1: TBS target identification on RS data

#### Participants

Twenty-nine young (range 20 – 35 years) right-handed (Oldfield, 1971) healthy individuals participated in this study as part of a bigger sample including different age groups and reported in previous research (Hermans et al., 2018; King et al., 2018; Levin et al., 2019; Monteiro et al., 2020; Zivari Adab et al., 2020). Participants had normal or corrected-to-normal vision, were not taking any psychoactive (e.g., anti-depressant or anxiety) medications, reported no known psychological, psychiatric or neurological disorders, and had no contra-indications for MRI. Of these 29 participants, three were excluded from the analyses: one due to RS fMRI data quality issues, one for co-registration failure (co-registration was not edited manually to keep the pre-processing procedure consistent across participants) and one for excessive movement during the RS scan (> 2.5 mm). Twenty-six participants (mean age: 25.48 ± 4.21 years, 11 females) were then included in the analyses. Note that the data were already available and that no sample size computation was thus performed when the current study was designed.

#### fMRI data acquisition and analysis

##### Acquisition

RS fMRI data were acquired with a Philips Achieva 3.0T MRI system equipped with a 32-channel head coil using an ascending gradient echo-planar imaging (EPI) pulse sequence for T2*-weighted images (TR = 2500 ms; TE = 30 ms; flip angle = 90°; 45 transverse slices; interslice gap = 0.25 mm; voxel size = 2.5 × 2.56 × 2.5 mm^3^; field of view = 200 × 200 × 123.5 mm^3^; matrix = 80 × 78; 162 dynamic scans plus 4 dummy scans discarded at the beginning of the sequence). Participants were instructed to keep their eyes open and to not think about anything in particular while the screen visible to them was turned to black (duration RS scan: 6 min 45 s). For each individual, a high-resolution T1-weighted structural image was also acquired with a magnetization-prepared rapid-acquisition gradient-echo (MPRAGE) sequence (TR/TE = 9.6/4.6 ms; voxel size = 1 × 1 × 1.2 mm^3^; field of view = 250 × 250 × 192 mm^3^; 160 coronal slices).

##### Pre-processing

RS fMRI data were preprocessed with a pipeline similar to King et al. (2018) using SPM12 (https://www.fil.ion.ucl.ac.uk/spm/software/spm12/; Wellcome Centre for Human Neuroimaging, London, UK) implemented in Matlab (The MathWorks, Natick, MA, USA). Each participant’s functional volumes were realigned to the first volume of the session using rigid body transformations and then slice time corrected. As a result of the realignment step, head motion was quantified and results showed limited movement during the RS scans. The absolute average ± SD of the maximum displacements across all resting state volumes and 3 planes of movement included in analyses was 0.70 ± 0.57 mm for linear translations and 0.79° ± 0.52° for rotations. Functional images were then co-registered to the anatomical image using a rigid body transformation optimized to maximize the normalized mutual information between the two images. The anatomical image was segmented into gray matter, white matter, cerebrospinal fluid (CSF), bone, soft tissue, and background. The mapping from participant to MNI space (Montreal Neurological Institute, http://www.bic.mni.mcgill.ca) was estimated from the anatomical image with the “unified segmentation” approach (Ashburner & Friston, 2005). The normalization parameters were subsequently applied to the individually co-registered functional volumes and the anatomical image.

##### Functional connectivity analyses

Functional connectivity (FC) analyses were conducted in Matlab with a pipeline similar to our previous research (King et al., 2018). Specifically, additional preprocessing steps were completed in order to remove variance from spurious sources before running connectivity analyses. First, to minimize the impact of motion on the correlations between voxels, volumes in which the scan-to-scan displacement exceeded 0.5 mm were removed and replaced via interpolation (mean: 5.96 ± 8.09%, range: 0 – 39.51% of acquired volumes discarded). Volumes were high-pass filtered using a 0.01 Hz cutoff. Average signals from the white matter and CSF were extracted using masks of these segments. Regression analyses were performed on the fMRI time-series, including the white matter and CSF signals (3 regressors each based on results from principal components analysis) as well as the 6-dimensional head motion realignment parameters, their squares, their derivatives, and the squared of the derivatives, as regressors. The resulting residuals were then low-pass filtered with a cutoff of 0.08 Hz. Data filtering served to minimize high-frequency noise that may be the result of cardiac and respiratory factors (Fox & Raichle, 2007; Fox et al., 2005). Finally, data were spatially smoothed with a Gaussian kernel of 6 mm full width half maximum (FWHM).

The goal of these connectivity analyses was to identify cortical regions reachable using TBS that were functionally and commonly connected to both the striatum and the hippocampus. To do so, we performed whole-brain FC analyses using the hippocampus and caudate nucleus (bilaterally, as defined anatomically according to the AAL brain atlas; Tzourio-Mazoyer et al., 2002) as seeds. Note that the striatal seed was restricted to the caudate nucleus, as this region exhibits functional and anatomical connectivity with the DLPFC (Albouy et al., 2012; Lehéricy et al., 2004), the TBS target region. For each individual and for each seed, the time-series across all voxels within the seed were averaged and Pearson correlation coefficients with all the voxels of the brain were computed. To ensure normality, each correlation coefficient was Fishers *r*-to-*z* transformed using the formula *z* = arctanh(*r*). Statistical analyses were performed on the z-values and were based on comparisons of the correlation coefficients to a value of 0. Statistical probabilities were considered significant if surviving the false discovery rate (FDR) method for multiple comparisons (*p*_FDR_ < .05). A conjunction analysis testing the “*Conjunction Null Hypothesis*” was performed between the hippocampal and striatal FDR-corrected connectivity Z-maps (hippocampus: Z ≥ 2.03, *p*_FDR_ < .05; caudate: Z ≥ 1.996, *p*_FDR_ < .05) using the easythresh_conj function (Nichols, 2007, http://www2.warwick.ac.uk/fac/sci/statistics/staff/academic-research/nichols/scripts/fsl/) rendering the conjunction map onto an average brain template provided by FSL (www.fmrib.ox.ac.uk/fsl, avg152T1) and thresholded at the highest Z score of both RSFC maps (Z = 2.03). The resulting statistical map showed any brain area that was significantly (*p* < .05) commonly connected to *both* seed regions at rest. Based on evidence reviewed above that (1) the DLPFC plays a pivotal role in the interaction between hippocampal and striatal systems during MSL (Albouy, King, et al., 2013) and that (2) repetitive TMS of the DLPFC can influence brain responses in these sub-cortical regions (e.g., Bilek et al., 2013; Ott et al., 2011), we constrained our TBS target search on the conjunction map to a mask including the middle and superior frontal segments of the AAL atlas (Tzourio-Mazoyer et al., 2002). The functional-data-driven DLPFC target was defined as the coordinate of the peak Z-value in the masked conjunction map.

#### Experiment 2: TBS and motor sequence learning

##### Participants

Twenty-one young (range: 19 - 26 years) right-handed (Oldfield, 1971) participants took part in this study. All participants had normal or corrected-to-normal vision, were nonsmokers, free of psychoactive (e.g., anti-depressant or -anxiety) medications, reported no known psychological, psychiatric or neurological disorders [including anxiety (Beck, Epstein, Brown, & Steer, 1988) and depression (Beck, Ward, Mendelson, Mock, & Erbaugh, 1961)], and had no contra-indications for MRI or TMS. Furthermore, none of the participants were considered musicians or professional typists. The quality and quantity of sleep during the month preceding the experiment was normal as assessed by the Pittsburgh Sleep Quality Index (Buysse, Reynolds, Monk, Berman, & Kupfer, 1989). Two participants were excluded because of incidental findings on the acquired imaging data. Nineteen participants were eventually included in the final analyses (see participants’ characteristics in Table 3). Due to technical problems, one experimental session (out of four) is missing for one participant. Behavioral, MRS and MRI data of two experimental sessions were excluded for another participant as he/she failed to appropriately perform the motor task (i.e., > 3 SD below the mean for accuracy). One session of another participant was excluded from the fMRI analyses due to excessive head motion (i.e., > 2 voxels). Motor Evoked Potential (MEP) data are missing for one participant. Consequently, behavioral, MEP, MRS and MRI analyses included 16 to 18 participants depending on the contrasts and conditions tested. Note that due to the multimodal nature of the present study, the choice of a specific outcome (among motor behavior, task-related activity, task- and resting-state-related connectivity, GABA levels) to perform sample size computation could be considered arbitrary. Consequently, our sample size estimation was based on previous studies that also sought to alter functional responses in sub-cortical areas via non-invasive brain stimulation applied to cortical targets. Previous research included on average 20 participants per group (Alkhasli et al., 2019; Bilek et al., 2013; Esslinger et al., 2014; Freedberg et al., 2019; Hanlon et al., 2016; Ott et al., 2011; van der Werf et al., 2010; Van Holstein et al., 2018; Wang et al., 2014), thus, this was our targeted sample size for Experiment 2.

### General experimental procedure

Participants were invited to complete five experimental sessions (one baseline and four TBS sessions) at the University Hospital of KU Leuven. All sessions occurred between 9am and 6pm. Moreover, all five sessions completed by each participant took place at approximately the same time of the day (± 2h) to minimize the influence of circadian phase variation on behavior (Smarr, Jennings, Driscoll, & Kriegsfeld, 2014), brain function (Muto et al., 2016) and brain excitability (de Beukelaar, Van Soom, Huber, & Wenderoth, 2016). TBS sessions were separated by at least 6 days (mean time between stimulation sessions: 7.9 ± 2.9 days) to avoid carry-over effects. Participants were instructed to have a good night of sleep before each experimental session and to avoid alcohol consumption the day before and the day of the experimental session. Sleep quality and quantity of the nights before each experimental session were assessed with the St. Mary’s Hospital Sleep Questionnaire (Ellis et al., 1981, see Table 3). Vigilance at the time of testing was assessed subjectively at the beginning of each session using the Stanford Sleepiness Scale (SSS; Maclean, Fekken, Saskin, & Knowles, 1992). Results related to the analyses of the sleep and vigilance data are reported in the Supplemental Material.

During the baseline MR session, a high-resolution T1-weighted image (to be subsequently used for neuronavigated TMS), RS functional data (to identify individual TBS targets, see below) as well as diffusion-weighted images (not reported in this manuscript) were acquired. Participants were also trained - for habituation purposes - on a random version of the serial reaction time task (see below). The session ended with a series of measures using the TMS equipment (determining the hot spot, resting and active motor thresholds, and corticospinal excitability through MEPs, see ‘TMS administration’ section). The next four experimental sessions were organized according to a stimulation (2 levels: intermittent TBS [iTBS] vs. continuous TBS [cTBS]) by task (2 levels: sequence [SEQ] vs. random [RND]) within-subject design (Figure 2; see below for details on the stimulation and task conditions). In each session, participants first underwent pre-TBS RS and MRS scans of the DLPFC and the hippocampus (see below for acquisition details) that were followed by T1-neuronavigated TBS applied to an individually-defined DLPFC target (see individual target identification below) outside the scanner. MEPs were measured pre- and post-stimulation as described below. Immediately following the end of the stimulation session, participants were placed in the MRI scanner where they were trained on the motor task while BOLD images were acquired (mean delay between start TBS and start task: 15.71 min, range 12 – 22; mean duration of the task training: 11.5 min, range 9.33 – 13.43). After task completion, post-TBS/task RS and MRS data of the DLPFC and hippocampus were acquired (intervals between TBS and post-TBS/task DLPFC and hippocampus MRS were 40.2 min, range: 36 – 46 and 51.85 min, range: 48 – 57, respectively; intervals between end of the task and post-TBS/task DLPFC and hippocampus MRS were 12.65 min, range: 12 – 15 and 24.29 min, range: 24 – 26, respectively). The order of the four experimental conditions [cTBS/SEQ (cSEQ), cTBS/RND (cRND), iTBS/SEQ (iSEQ), iTBS/RND (iRND)] was counterbalanced across participants.

### Serial reaction time task

An explicit bimanual version of the serial reaction time task (SRTT; Nissen & Bullemer, 1987) previously used in our group (King et al., 2019) that was coded and implemented with the Psychophysics Toolbox in Matlab (Brainard, 1997) was used in this study. Participants were lying in the scanner with a specialized MR-compatible keyboard placed on their lap. During the task, eight squares were presented on the screen via a mirror system above the participant’s head. Each square corresponded spatially to one of the eight keys on the keyboard and to one of eight fingers (excluding thumbs). The color of the outline of the squares alternated between red and green, indicating rest and practice blocks, respectively. After each rest block (15 s), the outlines of all squares changed from red to green, indicating that participants should prepare to perform the task. Subsequently, one of the eight squares was colored (i.e., filled) green, and participants were instructed to press the corresponding key with the corresponding finger as fast and as accurately as possible. As soon as a key was pressed, regardless of whether the response was correct or not, the next square in a sequence changed to green (response to stimulus interval = 0ms). Each block of practice included 48 key presses and each training session included 16 blocks. Depending on the specific experimental condition, the order in which the squares were filled green (and thus the order of finger movements) followed either a pseudorandom (RND) or a fixed, repeating sequential pattern (SEQ). During the sequence conditions, participants performed one of two eight-element sequences (whereby each of the eight fingers was pressed once in a sequence) that was repeated six times per block. The sequences were 4-7-3-8-6-2-5-1 and 7-2-8-4-1-6-3-5 with 1 representing the left little finger and 8 representing the right little finger, respectively. Note that due to experimental error, one participant was trained on sequences 4-7-3-8-6-2-5-1 and 2-6-1-5-8-3-7-4 and one participant was trained on 7-2-8-4-1-6-3-5 and 2-6-1-5-8-3-7-4. In the pseudorandom condition, there was no repeating sequence, but each key was pressed once every eight elements (i.e., no repeating elements); thus, each finger was also used six times per block. Participants were explicitly informed when the stimuli would follow a random pattern or a repeating sequential pattern but, in the latter case, they were not given any additional information such as what the pattern was or how many elements the sequence was composed of. Mean response time for correct responses (RT, reflecting performance speed) and percentage of correct responses (percentage correct, reflecting movement accuracy) were computed for each block. Data were analyzed using repeated measures analyses of variance (ANOVAs; α = .05) with stimulation (cTBS and iTBS), task (SEQ and RND) as well as block (1-16) as within-subject factors. Greenhouse-Geisser corrections were applied in case of violation of the sphericity assumption.

#### TMS administration

##### Individual target identification using baseline RS data

Individual TBS targets were identified using each participant’s RS data collected during the baseline session. RS fMRI data were acquired on a Philips Achieva 3.0T MRI system equipped with a 32-channel head coil using an ascending gradient EPI pulse sequence for T2*-weighted images (TR = 1000 ms; TE = 33 ms; multiband factor 3; flip angle = 80°; 42 transverse slices; interslice gap = 0.5 mm; voxel size = 2.14 × 2.18 × 3 mm^3^; field of view = 240 × 240 × 146.5 mm^3^; matrix= 112 × 110; 300 dynamic scans). Note that due to multiband capacity failure, the baseline RS data of one participant had different parameters: TR = 2500 ms; TE = 30 ms; flip angle = 90°; 45 transverse slices; slice thickness = 3 mm; interslice gap = 0.25 mm; voxel size 2.5 x 2.56 × 3 mm^3^; field of view = 200 × 200 × 146 mm^3^; matrix= 80 × 78; 162 dynamic scans. During data acquisition, a dark screen (i.e., no visual stimuli) was presented; participants were instructed to remain still, close their eyes and to not think of anything in particular for the duration of the scan (5 min). High-resolution T1-weighted structural images were acquired with a MPRAGE sequence (TR/TE = 9.6/4.6 ms; voxel size = 0.98 × 0.98 × 1.2 mm^3^; field of view = 250 × 250 × 228 mm^3^; 190 coronal slices). Four participants were scanned with a high-resolution T1-weighted structural MPRAGE sequence with the following parameters: TR/TE = 9.6/4.6 ms; voxel size = 0.98 × 0.98 × 1.2 mm^3^; field of view = 250 × 250 × 192 mm^3^; 160 coronal slices. RS data of each individual were preprocessed as described in Experiment 1. None of the subjects included in the analysis moved more than 1 voxel during the full duration of the scan. The absolute average ± SD of the maximum displacements across all resting state volumes and 3 planes of movement was 0.39 ± 0.16 mm for linear translations and 0.38° ± 0.24° for rotations. To minimize the impact of motion on the correlations between voxels, volumes in which the scan-to-scan displacement exceeded 0.5 mm were removed and replaced via interpolation (mean: 0.82 ± 1.04%, range: 0 – 3.33% of acquired volumes discarded). The individual’s TBS target was characterized using the same procedure as in Experiment 1 but applied at the individual level (i.e., conjunction between the individuals’ hippocampus and striatum RSFC maps) and using a 15-mm radius sphere mask centered on the group DLPFC coordinate identified in Experiment 1 rather than the AAL frontal mask for the target search (see Supplemental Table S6 for a list of individual TBS targets).

##### Theta-burst stimulation

TMS was applied, outside the MRI scanner, with a theta-burst stimulation (TBS) procedure (a burst of 3 pulses given at 50 Hz, repeated every 200 ms; Huang et al., 2005) on the individually-identified DLPFC target using a DuoMAG XT-100 rTMS stimulator (DEYMED Diagnostics s.r.o., Hronov, Czech Republic). Online spatial monitoring of the coil position was performed using neuronavigation (BrainSight, Rogue Research Inc, Montreal, Quebec, CA). We applied intermittent (iTBS, 2 s TBS trains repeated every 10 s for 190 s, 600 pulses) and continuous TBS (cTBS, 40 s uninterrupted train of TBS, 600 pulses) at 80% active motor threshold (MT, Huang et al., 2005). Active MT was characterized using single pulse stimulation of the M1 hotspot and motor evoked potentials (MEPs) measured with a belly-tendon EMG montage on the right flexor dorsal interosseous (FDI) muscle. Active MT was probed using a procedure similar to previous reports (Tambini, Nee, & D’Esposito, 2018; van Polanen, Rens, & Davare, 2019). Specifically, active MT was defined as the lowest intensity at which at least 5 out of 10 MEPs could be distinguished from background EMG during voluntary submaximal FDI contraction. During DLPFC TBS, the 70 mm DuoMAG butterfly coil was placed at a 45° angle with the handle pointing posteriorly. Subjects rested for 5 min post-TBS to not introduce any interfering effects of voluntary movements (Huang, Rothwell, Edwards, & Chen, 2008). Twenty-one MEPs at 120% resting MT were measured pre- and 5 min post-TBS (see Figure 2) as readout of corticospinal excitability (CSE) changes of M1. Resting MT was defined using single pulse stimulation of the M1 hotspot as the lowest intensity at which at least 5 out of 10 MEPs measured on the FDI were larger than 50 µV. The first MEP of each time point and session was discarded from analysis. For each participant and within each session, pre-TBS MEPs that were not within the range of the mean ± 3 SD were excluded (< 1% of all trials). For each experimental session, post-TBS MEPs were normalized to pre-TBS MEPs and a two-tailed paired t-test (α = .05) was performed to test for a stimulation effect (cTBS vs. iTBS; see Supplemental Figure S3). See Supplemental Material for results and discussion of the MEP data.

#### Magnetic Resonance Spectroscopy

##### Acquisition

In-vivo proton (^1^H) MRS (Mullins et al., 2014; Puts & Edden, 2012) was used to assess GABA+ levels in the DLPFC TBS target and the hippocampus. Before each MRS acquisition session, a low resolution T1-weighted structural image was acquired for MRS voxel positioning with a MPRAGE sequence (TR/TE = 9.6/4.6 ms; voxel size = 1.2 × 1.2 × 2.0 mm^3^; field of view = 250 × 250 × 222 mm^3^; 111 coronal slices). Lower-rather than higher-resolution scans were acquired due to time constraints but images showed sufficient quality to position the MRS voxel accurately. For each of the time points (pre-TBS and post-TBS/task) and for each condition, MRS data were acquired using the MEscher–GArwood Point RESolved Spectroscopy (MEGA-PRESS) sequence (Mescher, Merkle, Kirsch, Garwood, & Gruetter, 1998) over the individual DLPFC target (30 x 30 x 30 mm^3^ voxel) and the hippocampus (40 x 25 x 25 mm^3^ voxel) with parameters similar to previous research (Hermans et al., 2018; Maes et al., 2018): 320 averages, scan duration of 11 min, 14 ms editing pulses applied at an offset of 1.9 ppm in the ON experiment and 7.46 ppm in the OFF experiment, TR/TE = 2000/68 ms, 2-kHz spectral width, MOIST water suppression. Sixteen water-unsuppressed averages were acquired at each time point from the same voxel and interleaved to allow for real-time frequency correction (Edden et al., 2016), which is of special importance after fMRI scans (Harris et al., 2014). Scan parameters were identical for all MRS time points.

Before each MRS session, the TBS target was marked for each individual using a fiducial glycerin marker fixated on the participant’s head. The specific location on the skull was defined using the nudge tool of the Brainsight software that allows the projection of the individual MNI target coordinate onto the skull. All MRS voxels were positioned according to the MRS time point-specific, low-resolution T1 image. Specifically, the left DLPFC MRS voxel was positioned under this glycerin marker with one surface parallel to the cortical surface in the coronal and sagittal views (see Figure 4A for an example of voxel positioning and 4B for MRS spectra). The hippocampus voxel was positioned on the coronal view on the center of the left hippocampus and was aligned on the sagittal view parallel to the antero-posterior long axis. Note that we opted to not counterbalance the order of MRS voxel acquisitions and prioritized timing for the DLPFC voxel, as hippocampal MRS data analyses were considered as more exploratory. Therefore, the DLPFC voxel was always acquired before the hippocampus voxel so that the post-TBS/task measurement would be closer in time from the interventions (see Figure 2). Time constraints prevented us to acquire striatal MRS data as effects of TBS are thought to last on the order of 60 min (Huang et al., 2005). DLPFC and hippocampus voxel placement across sessions and participants are presented in the supplements (Supplemental Figure S1). Spatial overlap between sessions and participants was very high for the hippocampus voxel whereas consistency was lower for the DLPFC voxel as placement depended on the individually optimized TBS target.

##### Preprocessing and analyses

The Gannet software 3.0 toolkit (Edden, Puts, Harris, Barker, & Evans, 2014) was used for MRS data analysis similar to previous research in our group (Hermans et al., 2018; Maes et al., 2018). We corrected the individual frequency-domain spectra for frequency and phase using spectral registration in the time domain (Near et al., 2015). A 3 Hz exponential line broadening filter was applied subsequently. An edited difference spectrum was derived from the averaging and subtracting of individual ON and OFF spectra. The GABA signal from this difference spectrum was modelled at 3 ppm with a single Gaussian peak and a 5-parameter Gaussian model using the combined GABAGlx model. A Gaussian-Lorentzian model was used to fit the unsuppressed water signal that was used as the reference compound (Mikkelsen et al., 2019). Uncorrected GABA levels were quantified from the integrals of the modelled data. It is worth noting that this approach edits GABA as well as macromolecules at 3 ppm (Edden, Puts, & Barker, 2012; Rothman, Petroff, Behar, & Mattson, 1993) and thus GABA levels are reported as GABA+ (GABA plus macromolecules). The high-resolution T1-weighted image acquired during baseline was co-registered to the 8 (2 pre- and post-intervention time points x 4 conditions) low-resolution images using SPM12, so that the high-resolution structural image could be used for data processing for each MRS time point in each condition. MRS voxels were co-registered to the high-resolution T1-weighted image and were segmented into different tissue fractions (gray matter [GM], white matter [WM], and cerebrospinal fluid [CSF]) to adjust GABA+ levels for heterogeneity in voxel tissue composition. It was assumed that GABA+ levels are negligible in CSF and twice as high in GM relative to WM (Harris, Puts, & Edden, 2015) to compute tissue-corrected GABA+. Tissue-specific relaxation as well as water visibility values were also considered (Harris et al., 2015). Last, GABA+ levels were normalized to the average voxel composition in the sample (Harris et al., 2015). Therefore, the reported GABA+ values correspond to the “QuantNormTissCorrGABAiu” variable in Gannet 3.0, specified in institutional units [i.u.].

Due to low hippocampal MRS data quality, presumably due to difficulties associated with shimming in deep brain regions and participant movement between the low-resolution T1 (measured just before the RS, see Figure 2) and the hippocampal MRS scans, the fitting step as part of the Gannet pipeline failed in 15 out of 150 measurements during preprocessing. This resulted in 12 missing conditions, with a complete condition consisting of both the pre and post MRS time points for that particular experimental session (6 participants with 1 condition missing and 3 participants with 2 conditions missing). As too few measurements were left for appropriate statistical analyses of the hippocampal MRS data (only 10 participants with complete data sets), MRS analyses presented in this paper were limited to the DLPFC voxel.

Quality of the DLPFC MRS data was assessed by examining GABA signal-to-noise (SNR) ratio, fit error, and frequency offset. MRS voxel tissue fractions, quality metrics and corresponding statistical analyses to assess potential effects of MRS time point and experimental condition can be found in Supplemental Table S7. In sum, the averaged quality values are comparable to previous studies assessing GABA in the DLPFC (Hone-Blanchet et al., 2016; Mikkelsen, Loo, Puts, Edden, & Harris, 2018) as well as a recent, large multi-centre study that sought to provide quantitative benchmarks of quality metrics and GABA+ estimates, albeit from the parietal cortex (Mikkelsen et al., 2017, 2019). Importantly, these quality metrics did not differ between time points and experimental conditions.

For each experimental session, post-TBS/task GABA+ levels were normalized to pre-TBS GABA+ levels (*GABA+_pre_/GABA+_post_*, referred to as ΔGABA, see Supplemental Table S3 for raw data) and the data were analyzed using repeated measures analyses of variance (ANOVAs; α = .05) with stimulation (cTBS and iTBS) and task (SEQ and RND) as within-subject factors. Exploratory follow-up two-tailed paired t-tests (α = .05) were performed on all possible pairs. The individual normalized GABA+ data (ΔGABA) of each condition were also used as covariates for fMRI regression analyses (see details below).

#### Task-related fMRI data acquisition and analysis

##### Acquisition

Task-related fMRI data were acquired using an ascending gradient EPI pulse sequence for T2*-weighted images (TR = 2000 ms; TE = 29.8 ms; multiband factor 2; flip angle = 90°; 54 transverse slices; slice thickness = 2.5 mm; interslice gap = 0.2 mm; voxel size = 2.5 × 2.5 × 2.5 mm^3^; field of view = 210 × 210 × 145.6 mm^3^; matrix=84 × 82; 345.09 ± 22.37 dynamical scans).

##### Spatial pre-processing

Task-based functional volumes of each participant were realigned to the first image of each session and then realigned to the across-session mean functional image using rigid body transformations. The mean functional image was co-registered to the high-resolution T1-weighted anatomical image using a rigid body transformation optimized to maximize the normalized mutual information between the two images. The resulting co-registration parameters were then applied to the realigned functional images. The structural image was segmented into gray matter, white matter, cerebrospinal fluid (CSF), bone, soft tissue, and background. We created an average subject-based template using DARTEL in SPM12, registered to the Montreal Neurological Institute (MNI) space. All functional and anatomical images were then normalized to the resulting template. Functional images were spatially smoothed using an isotropic 8 mm full-width at half-maximum (FWHM) Gaussian kernel.

##### Activation analyses

The analysis of task-based fMRI data, based on a summary statistics approach, was conducted in 2 serial steps accounting for intra-individual (fixed effects) and inter-individual (random effects) variance, respectively. Changes in brain regional responses were estimated for each participant with a model including responses to the motor task and its linear modulation by performance speed (mean RT on correct button presses per block) in each session (cSEQ, cRND, iSEQ and iRND). Performance speed, rather than accuracy, was chosen as a parametric modulator because performance accuracy remained stable during practice (see results section) and was therefore not modulated by task practice. The 15-second rest blocks occurring between each block of motor practice served as the baseline condition modeled implicitly in the block design. These regressors consisted of box cars convolved with the canonical hemodynamic response function. Movement parameters derived from realignment as well as erroneous key presses were included as covariates of no interest. Movements were minimal during scanning; only the data of one session in one participant were excluded for excessive movement (> 2 voxels; note that for another participant, the last 46 scans of one session were excluded from analyses because of movements but the truncated session was kept in the analyses). The average ± SD translation and rotation across axis and sessions was: 1.07 ± 0.62 mm and 1.10 ± 0.61° (maximum absolute movement in translation = 3.7 mm and in rotation = 2.9°). High-pass filtering was implemented in the design matrix using a cutoff period of 128s to remove slow drifts from the time series. Serial correlations in the fMRI signal were estimated using an autoregressive (order 1) plus white noise model and a restricted maximum likelihood (ReML) algorithm.

Linear contrasts tested the main effect of practice and its linear modulation by performance speed in each session as well as between sessions. Contrasts testing for the stimulation by task interaction [(iTBS vs. cTBS) x (SEQ vs. RND)] and the stimulation effect within each task condition [iSEQ vs. cSEQ] and [iRND vs. cRND] were generated at the individual level. To examine whether the dynamics of brain responses were influenced by stimulation conditions, contrasts tested for the stimulation effect on the modulation regressors. As performance levels remained – as expected - constant in the random conditions (see results), this set of analyses focused on the sequence conditions only [iSEQ_mod_ vs. cSEQ_mod_]. Additional contrasts presented in the supplemental data tested for the main effect of practice across task conditions [SEQ+RND] (see Supplemental Table S8) as well as the modulation effect across stimulation conditions within the sequence task [iSEQ_mod_+cSEQ_mod_] (see Supplemental Table S4). The resulting contrast images were further spatially smoothed (Gaussian kernel 6 mm FWHM) and were entered in a second level analysis for statistical inference at the group level (one sample t-tests), corresponding to a random effects model accounting for inter-subject variance.

To assess the relationship between any effect highlighted in the contrasts described above and the pre- to post-intervention changes in GABA+ levels (referred to as ΔGABA), we performed regression analyses at the second level using one sample t-test with multiple covariates. Specifically, we regressed the individual contrast images testing for the stimulation by task interaction [(iSEQ - iRND) - (cSEQ - cRND)] against individual ΔGABA measured in the four conditions (4 covariates). The multiple regression therefore tested whether stimulation by task-related activity patterns correlated with stimulation by task-related changes in GABA levels in the DLPFC [(ΔGABA_iSEQ_ - ΔGABA_iRND_) – (ΔGABA_cSEQ_ - ΔGABA_cRND_)]. A separate multiple regression analysis tested whether the stimulation effect on dynamical activity within the SEQ task condition [iSEQ_mod_ vs. cSEQ_mod_] correlated with the stimulation effect on ΔGABA in the corresponding conditions [ΔGABA_iSEQ_ vs. ΔGABA_cSEQ_]. In these regression analyses, any significant brain response is differently related to ΔGABA between stimulation (or stimulation by task; for the interaction contrast) conditions.

##### Functional connectivity analyses

Psychophysiological interaction (PPI) analyses were computed to test the functional connectivity of the individual DLPFC targets and subcortical a priori regions of interest (i.e. the striatum and the hippocampus) highlighted by the activation-based contrasts. Seed coordinates for the DLPFC connectivity analyses consisted of the individual TBS targets as identified with the RS pipeline (see above). Note that the group, rather than the individual, target was used in two participants as their individual coordinates were located close to the cortex’s edge which did not allow the extraction of enough seed signal (see procedure below). Two putamen, but no hippocampal, seed regions were identified based on activation analyses. PPI analyses were performed using the peak coordinate of the two significant putamen clusters highlighted in the group level activation maps (iSEQ_mod_+cSEQ_mod_, see Supplemental Table S4; [24 12 4 mm] and [-16 6 −6 mm]). For each participant, experimental session and seed region of interest, the first eigenvariate of the signal was extracted using Singular Value Decomposition of the time series across the voxels included in a 10 mm radius sphere centered around the seed of interest. A new linear model was generated at the individual level, using three regressors for each experimental session. The first regressor corresponded to the BOLD activity in the reference area. The second regressor represented the practice of the learned sequence or the practice of the learned sequence modulated by performance speed. The third regressor represented the interaction of interest between the first (physiological) and the second (psychological) regressors. To build this regressor, the underlying neuronal activity was first estimated by a parametric empirical Bayes formulation, combined with the psychological factor, and subsequently convolved with the hemodynamic response function (Gitelman, Penny, Ashburner, & Friston, 2003). The design matrix also included movement parameters. A significant PPI indicated a change in the regression coefficients (i.e. a change in the strength of the functional interaction) between any reported brain area and the reference region, related to the practice of the task or to the change in performance speed during the practice of the task. Linear contrasts testing the stimulation by task interaction [(iTBS vs. cTBS) x (SEQ vs. RND)] as well as the main effect of stimulation on modulation within SEQ conditions [iSEQ_mod_ vs. cSEQ_mod_] were generated at the individual level. The resulting contrast images were further spatially smoothed (Gaussian kernel 6 mm FWHM) and were entered in a second level analysis for statistical inference at the group level (one sample t-tests), corresponding to a random effects model accounting for inter-subject variance. Furthermore, we assessed the relationship between DLPFC connectivity patterns and ΔGABA levels in the DLPFC with regression analyses at the second level using one sample t-test with multiple covariates. As no significant responses were observed for the DLPFC connectivity analyses on the interaction contrast (see results), regression analyses were only performed on the DLPFC PPI analyses testing for the stimulation effect within SEQ conditions. Specifically, we regressed the individual contrast images testing for the difference in dynamical connectivity between the two SEQ conditions [iSEQ_mod_ vs. cSEQ_mod_] against the ΔGABA in these two conditions [ΔGABA_iSEQ_ vs. ΔGABA_cSEQ_]. In these analyses, any significant brain response shows connectivity patterns with the DLPFC during sequence learning that are differently related to the change in DLPFC GABA between stimulation conditions.

##### Statistical inferences

The set of voxel values resulting from each analysis described above (activation and functional connectivity) constituted maps of the t statistics [SPM(T)], thresholded at *p* < .005 (uncorrected for multiple comparisons). Statistical inferences were performed at a threshold of *p* < .05 after family-wise error (FWE) correction for multiple comparisons over small volume (SVC, 10 mm radius) located in structures of interest reported by published work on motor learning (see Supplemental Material).

## Acknowledgements

This work was supported by the Belgian Research Foundation Flanders (FWO; G099516N) and internal funds from KU Leuven. GA also received support from FWO (G0D7918N, G0B1419N, 1524218N) and Excellence of Science (EOS, 30446199, MEMODYN, with SPS and DM). MAG, ND and MPV received salary support from these grants. MAG is funded by a predoctoral fellowship from FWO (1141320N). Financial support for author BRK was provided by the European Union’s Horizon 2020 research and innovation program under the Marie Skłodowska-Curie grant agreement (703490) and a postdoctoral fellowship from FWO (132635). This study applies tools developed under National Institutes of Health (NIH) Grants R01-EB-016089, R01-023963, and P41-EB015909; RAEE also receives salary support from these grants. NAJP receives salary support from NIH Grant R00-MH-107719. EMR received salary support from the Air Force Office of Scientific Research (AFOSR, Virginia, USA; FA9550-16-1-0191). We wish to thank Kaat van Rooij and Michelle Roussard for assistance with data collection.

## Author contributions

GA, EMR, BRK, MD, MAG designed the experiments. MAG conducted the experiments in collaboration with ND, MVP, and GA. GA, MAG, BRK, ND, NAJP, RAEE, KLC, MD, SPS and DM contributed to acquisition and analytic tools. MAG analysed the data in collaboration with GA & BRK. MAG and GA drafted the manuscript. All authors revised the manuscript.

## Data availability statements

The ethical approval granted by the local ethics committee does not permit the publication of raw data online. If readers would like to access these data, additional ethical approval (on an individual user and purpose basis) will be required. The authors are happy to support such ethical approval applications.

## Supplemental Material

### Supplemental Methods

#### Coordinates of areas of interest used for spherical small volume corrections

<u>Frontal:</u> Superior frontal cortex −36 50 22 mm, 24 44 32 mm, −32 46 32 mm, ±30 18 58 mm, ±26 40 30 mm (Albouy et al., 2008); 32 54 22 mm (Oishi et al., 2005); 50 18 44 mm (Albouy et al., 2013); Inferior frontal gyrus ±36 46 −4 mm, ±36 16 24 mm (Albouy et al., 2008); Middle frontal cortex −44 40 26 mm, 36 50 22 mm (Albouy et al., 2008); −36 0 42 mm (Albouy et al., 2015); ±30 8 63 mm (Gheysen et al., 2016); Medial prefrontal cortex 30 28 34 mm (Bischoff-Grethe, Goedert, Willingham, & Grafton, 2004); 6 60 2 mm (Sterpenich et al., 2007); ±26 48 2 mm (Fernández-Seara, Aznárez-Sanado, Mengual, Loayza, & Pastor, 2009); 10 48 −8 mm (Albouy et al., 2015)

<u>Parietal:</u> Superior parietal cortex ±12 −62 62 mm (Penhune & Doyon, 2005); 11 −52 71 mm (Lungu et al., 2014); Inferior parietal cortex 36 −52 39 (= 35 −55 42 mm in MNI) (Grafton, Hazeltine, & Ivry, 1995); Intraparietal sulcus ±34 −66 36 mm, ±38 −64 54 mm (Sakai, Ramnani, & Passingham, 2002); −48 −60 48 mm, −52 −46 38 mm (Albouy et al., 2015); 30 −54 70 mm, ±22 −64 44 mm, ±48 −50 60 mm (Albouy et al., 2008); 48 −66 30 mm (Penhune & Doyon, 2002); Supramarginal gyrus −52 −40 28 mm (Albouy et al., 2008); Precuneus ±4 −58 66 mm (Albouy et al., 2015)

<u>Motor/central:</u> Precentral gyrus −54 0 44 mm, 48 8 30 mm (Albouy et al., 2008); Postcentral gyrus 22 −44 70 mm (Albouy et al., 2015); ±44 −20 37 (Gheysen et al., 2016); Premotor cortex ±20 −4 46 mm, 12 −14 64 mm (Penhune & Doyon, 2005); Premotor cortex/Supplementary motor area ±10 0 72 mm (Penhune & Doyon, 2005); Supplementary motor area 4 −8 54 mm (Penhune & Doyon, 2005); M1/SM1 50 −18 42 mm (Penhune & Doyon, 2005)

<u>Para-/Hippocampal:</u> Parahippocampal gyrus ±26 −4 −38 mm (Penhune & Doyon, 2002); Parahippocampal gyrus/ Hippocampus head −33 −18 −27 mm (Schendan, Searl, Melrose, & Stern, 2003); Posterior hippocampus 26 −34 −6 mm (Albouy et al., 2008)

<u>Striatal:</u> Putamen ±28 −14 −8 mm, ±28 8 18 (Albouy et al., 2008); Ventral putamen −14 10 −10 mm (Albouy et al., 2008); Dorsal putamen ±27 6 −6 mm (Schendan et al., 2003); Caudate 9 15 −3 (Schendan et al., 2003)

<u>Cerebellar:</u> Cerebellar Lobule VI 30 −40 −24 mm (Albouy et al., 2015); Cerebellar Lobule VIIIB ±26 −44 − 42 mm (Penhune & Doyon, 2002); Cerebellar Lobule IX −6 −58 −44 mm (Penhune & Doyon, 2002); Cerebellar hemisphere ±18 −44 −18 mm (Fischer, 2005); ±26 −44 −42 mm (Penhune & Doyon, 2002); ±18 −72 −36 mm (Penhune & Doyon, 2005)

<u>Insular:</u> Anterior insula ±18 12 −14 mm (Penhune & Doyon, 2005); Insular gyrus 44 26 6 mm, −34 20 4 mm, 42 −4 −6 mm (Albouy et al., 2008)

<u>Cingulate:</u> Anterior cingulate cortex ±14 21 41 mm (Grafton, Hazeltine, & Ivry, 1998); ±12 30 −3 (Grafton et al., 1995); Anterior cingulate hand movement region −2 32 2 mm, 4 20 38 mm (Amiez & Petrides, 2014)

<u>Thalamus:</u> Thalamus ±10 −10 2 mm (Albouy et al., 2008)

### Supplemental Results

#### Sleep Quality & Quantity, Vigilance

Sleep quality and quantity (see Supplemental Table S5) during the night preceding each experimental session was assessed with the St. Mary’s Hospital Sleep Questionnaire (Ellis et al., 1981). A 2×2 repeated measures ANOVA with stimulation (iTBS vs. cTBS) and task (SEQ vs. RND) as within-subject factors revealed no significant main effect of stimulation (F_(1,16)_ = .027, *p* = .872), main effect of task (F_(1,16)_ = 1.15, *p* = .3) or interaction (F_(1,16)_ = 1.739, *p* = .206) for sleep quality. Similarly, there was no significant main effect of stimulation (F_(1,17)_ = .004, *p* = .949), main effect of task (F_(1,17)_ = .615, *p* = .444) or stimulation by task interaction (F_(1,17)_ = 3.647, *p* = .073) for sleep quantity. Note that sleep quality is missing for one participant in one condition (cSEQ) and both quality and quantity are missing for one condition (cRND) in another participant.

A 2×2 repeated measures ANOVA with stimulation (iTBS vs. cTBS) and task (SEQ vs. RND) as within-subject factors on subjective vigilance (see Supplemental Table S5) assessed with the Stanford Sleepiness Scale (SSS; Maclean, Fekken, Saskin, & Knowles, 1992) revealed no main effect of stimulation (F_(1,12)_ = .081, *p* = .781), but a significant main effect of task (F_(1,12)_ = 6.169, *p* = .029) and a significant stimulation by task interaction (F_(1,12)_ = 13.559, *p* = .003). Unexpectedly, participants indicated higher subjective vigilance (i.e., less sleepiness) in iRND compared to iSEQ (t_(14)_ = 2.416, *p* = .03) and cRND (t_(15)_ = 2.535, *p* = .023). It should be noted, however, that there were no significant correlations between the subjective vigilance ratings and the average performance speed in the corresponding conditions (cSEQ: r = −.033, *p* = .905; cRND: r = −.194, *p* = .44; iSEQ: r = −.24, *p* = .353; iRND: r = .014, *p* = .959). Thus, these condition differences did not appear to influence motor behavior. It is also worth mentioning that SSS data are missing in two conditions (iSEQ, cSEQ) for one participant and in one condition for five participants (1x iSEQ, 2x iRND, 1x cSEQ, 1x cRND).

Altogether, these results indicate that there were no differences in sleep quantity and quality and that differences in subjective vigilance did not influence performance speed.

#### Quality metrics related to DLPFC MRS of GABA

MRS voxel tissue fractions as well as data quality metrics for each MRS time point and condition are detailed in Supplementary Table S7. Overall, the averaged quality values are comparable to previous studies assessing GABA in the DLPFC (Hone-Blanchet, Edden, & Fecteau, 2016; Mikkelsen, Loo, Puts, Edden, & Harris, 2018) as well as a recent, large multi-centre study that sought to provide quantitative benchmarks of quality metrics and GABA estimates, albeit from the parietal cortex (Mikkelsen et al., 2017, 2019). There were no group differences nor any significant interactions between time point, stimulation and motor task for GABA signal-to-noise ratio (SNR), GABA fit error, average center of the frequency offset or WM tissue fractions (statistical results reported in Supplemental Table S7). However, the analyses yielded a significant main effect of stimulation for GM tissue fractions (F_(1,16)_ = 4.672, *p* = .046). Specifically, GM tissue fractions were unexpectedly higher in the cTBS compared to the iTBS conditions (t_(16)_ = 2.161, *p* = .046). There was also a significant time point x stimulation x task interaction effect for CSF tissue fractions (F_(1,16)_ = 5.263, *p* = .036), a result that was driven by stimulation x time point effect within the SEQ task condition (F_(1,16)_ = 5.611, *p* = .027). Specifically, while CSF fractions within iSEQ stayed rather stable across the two time points, CSF fraction increased in the cSEQ conditions across time points (t_(16)_ = 2.370, *p* = .031). Although these differences in voxel composition were unexpected, it is unlikely they exerted any influence on the results presented in the main text. First, our measure of GABA+ levels takes voxel composition into account, so any variations are considered in the output measure. Second, exploratory correlation analyses revealed that GABA+ levels were not related to CSF tissue fraction in the cSEQ condition (r = .156, *p* = .379), with the GM tissue fraction in the cTBS conditions (r = − .012, *p* = .925) nor with the GM tissue fraction in the iTBS conditions (r = − .018, *p* = .886).

As a final point of emphasis, MRS data are missing for one condition (cRND) in one participant due to technical problems and two conditions (cSEQ, iRND) for another participant due to outlier behavior on accuracy.

### MEPs

A two-tailed paired t-test on the post-TBS MEPs normalized to pre-TBS MEPs showed that MEP amplitude tended to be higher after iTBS as compared to cTBS; however, the difference only exhibited a trend for statistical significance (t_(17)_ = 1.921, *p* = .072; see Supplemental Figure S3). These results suggest that iTBS and cTBS of the DLPFC did not differently modulate corticospinal excitability (CSE) of M1.

### Supplemental Discussion

#### DLPFC stimulation did not modulate corticospinal excitability

In the current study, DLPFC stimulation did not induce any significant changes in CSE measured on M1. Based on evidence showing inhibitory interactions between the DLPFC and M1 at rest (Cao et al., 2018; Civardi, Cantello, Asselman, & Rothwell, 2001; Rollnik, Schubert, & Dengler, 2000), one could have expected DLPFC cTBS and iTBS to respectively increase and decrease CSE. While the effect of frontal stimulation on CSE has not typically been tested or reported in the majority of previous research (Alkhasli, Sakreida, Mottaghy, & Binkofski, 2019; Bilek et al., 2013; Chung et al., 2017; Dayan, Herszage, Laor-Maayany, Sharon, & Censor, 2018; Galea, Albert, Ditye, & Miall, 2010; Robertson, Tormos, Maeda, & Pascual-Leone, 2001), studies in which this measure is provided show discrepant results. On the one hand, and in line with our results, several studies showed no significant modulation of CSE measured with MEPs following repetitive TMS (rTMS) of the DLPFC (Do et al., 2018; Fierro et al., 2010; Rens et al., 2020). On the other hand, other studies showed modulation of MEP magnitude after DLPFC stimulation that are in line with the above-described inhibitory DLPFC-M1 relationship (Cao et al., 2018; Rollnik et al., 2000). For example, Rollnik et al. (2000) applied facilitatory 5 Hz rTMS to the DLPFC and showed that MEP area decreased during DLPFC stimulation. In agreement with these findings, Cao et al. (2018) measured MEPs at different time points post DLPFC stimulation (from immediately to up to 60 min post-stimulation) and showed that continuous and intermittent TBS induced an increase and decrease in MEP amplitude, respectively. While this has never been the subject of a systematic investigation, it is possible that discrepancies between studies are explained by differences in stimulation type, timing of the MEP measurements and target definition.

## Supplemental Figures

**Supplemental Figure S1.**
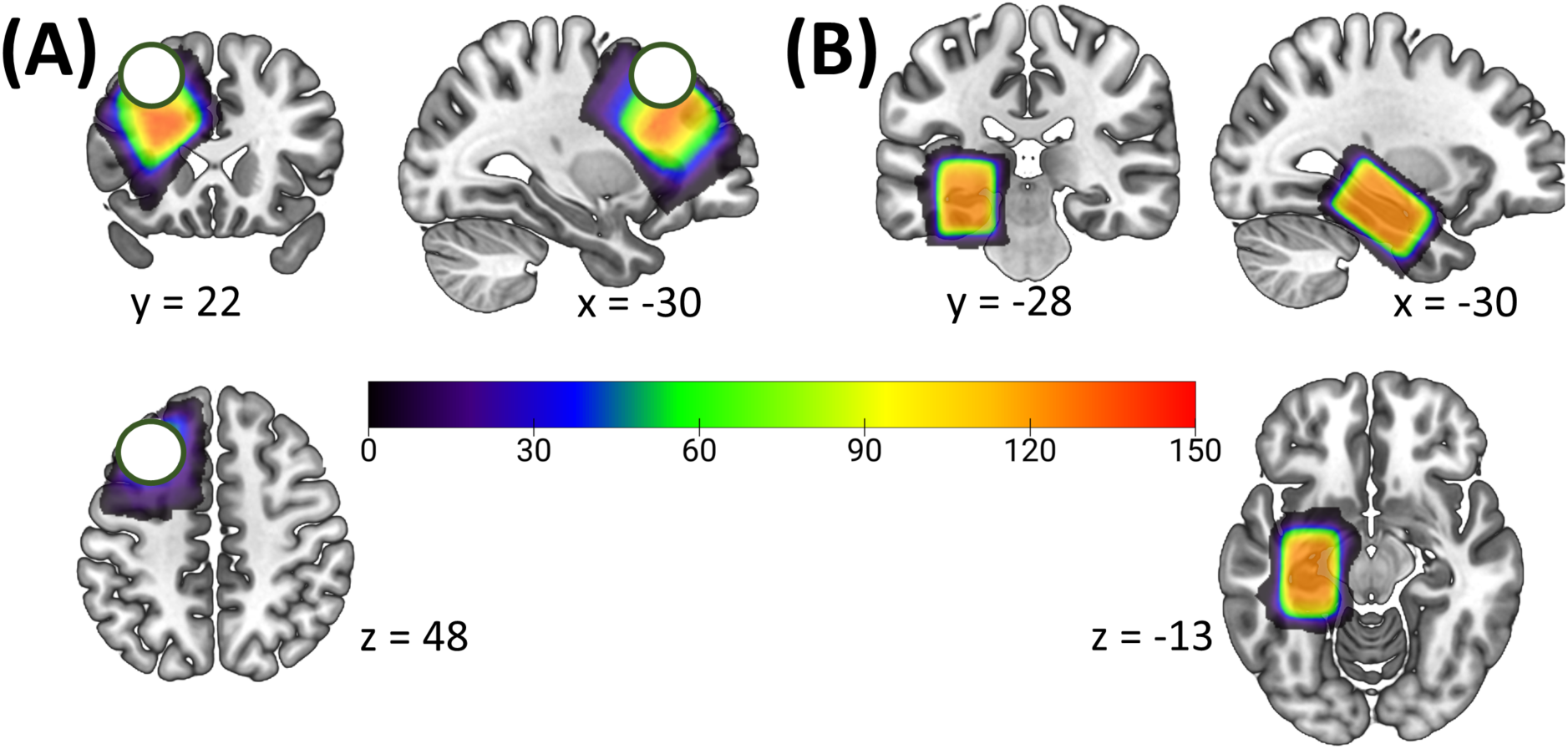
Heatmaps representing the spatial overlap of voxel placement for the (A) DLPFC and (B) hippocampal MRS voxels across all conditions and time points, overlaid on a template image. Color bars represent the number of overlapping voxels. The 15-mm radius search sphere for the individual targeting pipeline in Experiment 2 is indicated with a white sphere on (A). While consistency in voxel placement was high across individuals and time points for the hippocampal voxel, it was lower for the DLPFC due to the individualized approach used for TMS targeting. DLPFC: dorsolateral prefrontal cortex, MRS: magnetic resonance spectroscopy.

**Supplemental Figure S2.**
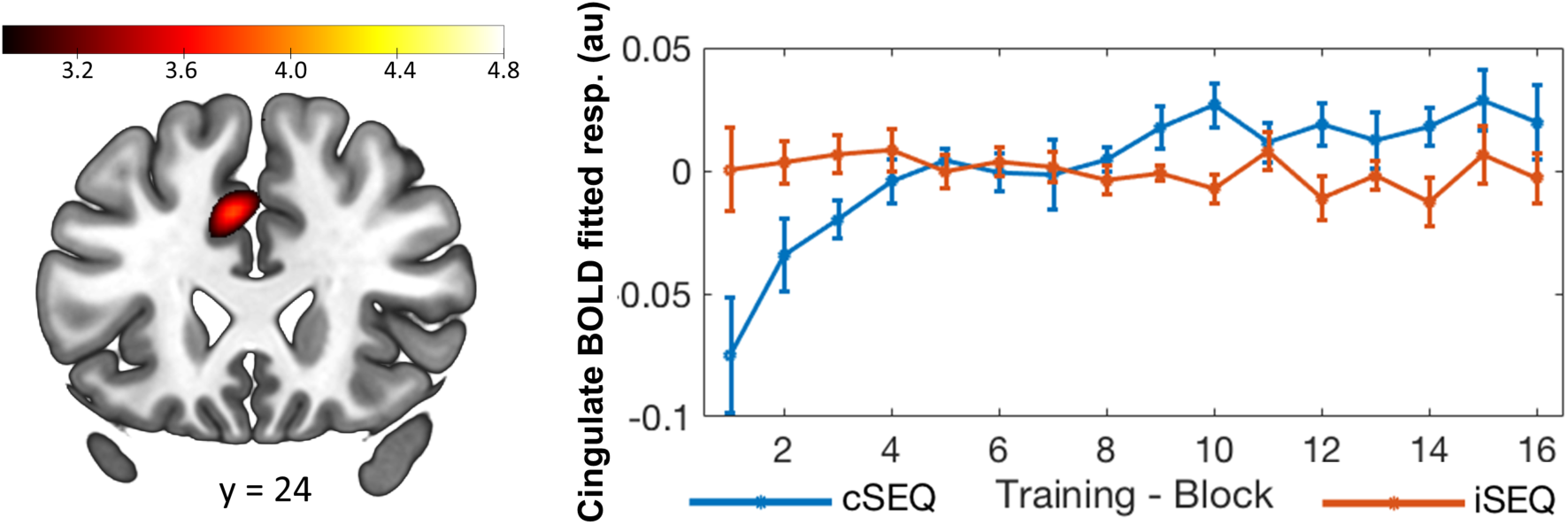
The activity in the cingulate motor area (left panel) during sequence learning (SEQ) was differently modulated by performance depending on the stimulation condition. This effect was driven by a practice-related increase in activity in cSEQ. Activations maps are displayed on a T1-weighted template image with a threshold of *p* < .005 uncorrected. Color bars represent T values. Error bars indicate SEM. au: arbitrary units, resp.: response, c: continuous, i: intermittent.

**Supplemental Figure S3.**
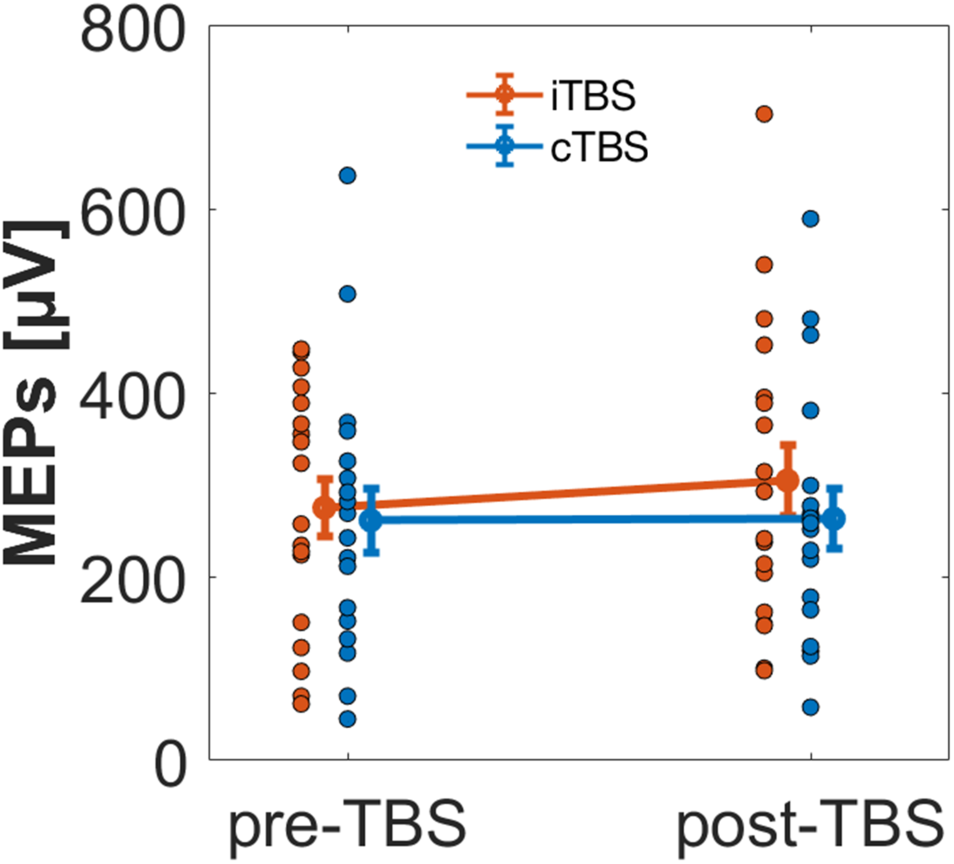
Raw MEP data averaged across task conditions within each stimulation condition and session (pre- and post-TBS). MEP amplitude tended to be higher after iTBS as compared to cTBS. MEP = motor evoked potential, iTBS = intermittent theta burst stimulation; cTBS = continuous theta burst stimulation.

## Supplemental Tables

**Supplemental Table S1:** Resting-state functional connectivity (RSFC) results from Experiment 1

| Area | x mm | y mm | z mm | k voxels | Z |
| --- | --- | --- | --- | --- | --- |
| ***Hippocampi RSFC*** |  |  |  |  |  |
| Hippocampus | -22 | -22 | -14 | 49253 | 10.9 |
| Parahippocampal gyrus |  |  |  |  |  |
| Fusiform gyrus |  |  |  |  |  |
| Amygdala |  |  |  |  |  |
| Olfactory cortex |  |  |  |  |  |
| Caudate |  |  |  |  |  |
| Cingulum |  |  |  |  |  |
| Precuneus |  |  |  |  |  |
| Cuneus |  |  |  |  |  |
| Superior and middle frontal gyrus |  |  |  |  |  |
| Supplementary motor area |  |  |  |  |  |
| Insula |  |  |  |  |  |
| Cerebellum | 10 | -54 | -40 | 510 | 4.54 |
|  | 10 | -88 | -38 | 115 | 3.49 |
|  | 44 | -72 | -38 | 114 | 3.8 |
|  | -52 | -46 | -44 | 39 | 2.77 |
|  | 60 | -48 | -34 | 35 | 2.83 |
| Inferior frontal gyrus | 32 | 36 | -10 | 245 | 4.22 |
|  | -56 | 26 | 16 | 56 | 2.95 |
|  | 52 | 26 | 0 | 35 | 2.6 |
| Supplementary motor area | -16 | -2 | 68 | 78 | 3.42 |
|  | -2 | 22 | 68 | 43 | 2.84 |
| Superior parietal gyrus | 22 | -58 | 56 | 34 | 3.14 |
| Caudate nucleus | 10 | 26 | 4 | 32 | 3.3 |
| Precentral sulcus | 58 | 6 | 28 | 30 | 2.83 |
|  | -58 | 6 | 34 | 20 | 2.76 |
| ***Caudate Nuclei RSFC*** |  |  |  |  |  |
| Caudate nucleus | 10 | 6 | 6 | 53344 | 9.64 |
| Putamen |  |  |  |  |  |
| Thalamus |  |  |  |  |  |
| Pallidum |  |  |  |  |  |
| Amygdala |  |  |  |  |  |
| Insula |  |  |  |  |  |
| Temporal pole |  |  |  |  |  |
| Inferior, medial and superior frontal gyrus |  |  |  |  |  |
| Superior temporal gyrus |  |  |  |  |  |
| Cingulum |  |  |  |  |  |
| Supplementary motor area |  |  |  |  |  |
| Inferior parietal gyrus | 50 | -38 | 56 | 1458 | 3.56 |
| Angular gyrus |  |  |  |  |  |
| Postcentral gyrus |  |  |  |  |  |
| Supramarginal gyrus |  |  |  |  |  |
| Inferior occipital gyrus | 30 | -92 | -12 | 278 | 3.56 |
| Middle occipital gyrus | -20 | -106 | 2 | 268 | 3.26 |
| Temporal Pole | -44 | 14 | -32 | 78 | 4.21 |
| Inferior and middle temporal gyrus |  |  |  |  |  |
| Inferior temporal gyrus | 42 | -6 | -40 | 69 | 3.25 |
| Fusiform gyrus |  |  |  |  |  |
| Inferior temporal gyrus | -68 | -40 | -20 | 48 | 3.09 |
| Middle temporal gyrus | 58 | -68 | 16 | 44 | 2.62 |
| Postcentral gyrus | 54 | -10 | 26 | 39 | 3.08 |
|  | 28 | -28 | 58 | 38 | 3.27 |
| Paracentral lobule | -10 | -34 | 64 | 36 | 2.65 |
| Precentral gyrus | 20 | -24 | 76 | 33 | 2.44 |
| Insula | 36 | -14 | 14 | 30 | 3.8 |
| Superior frontal gyrus | 22 | 10 | 66 | 27 | 3.02 |
| Calcarine fissure | -8 | -104 | -4 | 26 | 2.59 |
| Middle and superior occipital gyrus | 26 | -98 | 10 | 25 | 2.61 |

*Brain activations significant after false discovery rate (FDR) correction ( $p_{corr} < .05$ ) for multiple comparisons over the entire brain volume are reported here.*

**Supplemental Table S2:**
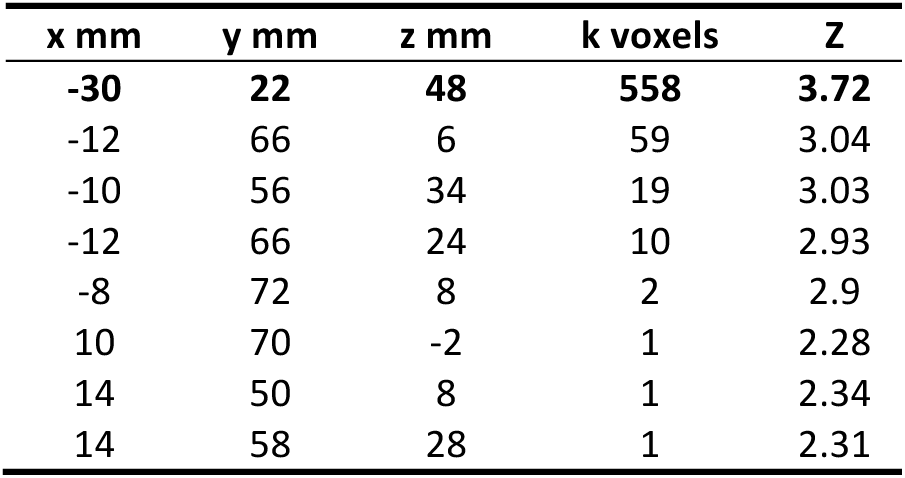
List of prefrontal clusters identified in the conjunction analysis between hippocampal and caudate resting-state functional connectivity maps. The peak cluster highlighted in bold was used as the center of a search sphere on individual’s conjunction maps in Experiment 2.

**Supplemental Table S3:** GABA+ values and relative change in GABA+ from pre-TBS to post-TBS/task in the DLPFC voxel

|  |  | cSEQ (n=18) | cRND (n=18) | iSEQ (n=19) | iRND (n=18) |
| --- | --- | --- | --- | --- | --- |
| **GABA+ [i.u.]** | pre | 2.57±.29 | 2.61±.15 | 2.56±.022 | 2.47±.27 |
|  | post | 2.64±.23 | 2.66±.25 | 2.48±.18 | 2.58±.17 |
| **Δ GABA** |  | 1.03±.12 | 1.02±.11 | .97±.11 | 1.06±.12 |

All GABA+ values are QuantNormTissCorrGABA as provided in Gannet 3.0. ΔGABA reflects the normalized change from pre to post time points computed as $GABA^{+}_{pre-TBS} / GABA^{+}_{post-TBS/task}$ . Participants completed 4 experimental conditions [cTBS/SEQ (cSEQ), cTBS/RND (cRND), iTBS/SEQ (iSEQ), iTBS/RND (iRND)]. GABA = gamma-aminobutyric acid, i.u. = institutional unit, SEQ = sequential motor task, RND = random motor task, iTBS = intermittent theta burst stimulation, cTBS = continuous theta burst stimulation, DLPFC = dorsolateral prefrontal cortex.

**Supplemental Table S4:** Modulation of brain responses by performance speed during practice of the sequential motor task (iSEQ_mod_+ cSEQ_mod_)

| Area | x mm | y mm | z mm | k voxels | T |
| --- | --- | --- | --- | --- | --- |
| ***Decrease in activity as a function of performance improvement*** |  |  |  |  |  |
| Cuneus, occipital cortex | -8 | -94 | 22 | 628 | 4.97 |
|  | -12 | -86 | 42 |  | 4.48 |
|  | 10 | -90 | 30 |  | 3.65 |
|  | 16 | -84 | 44 | 3 | 2.96 |
| Precuneus | -18 | -44 | 52 | 59 | 3.87 |
| Rolandic operculum | -52 | -6 | 10 | 24 | 3.31 |
| Insula | 34 | -26 | 24 | 3 | 3.28 |
|  | -38 | 0 | 16 | 14 | 3.21 |
| Parahippocampus | 32 | -14 | -32 | 8 | 3.21 |
| Superior temporal sulcus | 64 | -54 | 24 | 9 | 3.16 |
|  | 44 | -78 | 18 | 2 | 2.97 |
| Amygdala | 32 | 0 | -26 | 7 | 3.03 |
| Lateral fissure | -42 | -30 | 22 | 4 | 2.96 |
| ***Increase in activity as a function of performance improvement*** |  |  |  |  |  |
| Fusiform gyrus | 38 | -46 | -8 | 97 | 5.71 |
| Putamen | 24 | 12 | 4 | 660 | 5.03 |
|  | -16 | 6 | -6 |  | 4.66 |
| Frontal cortex | -14 | 46 | -10 | 1113 | 4.74 |
|  | 12 | 38 | -12 | 18 | 3.49 |
|  | 18 | -6 | 54 | 53 | 3.47 |
| Cerebellum | 20 | -52 | -34 | 277 | 4.09 |
|  | 34 | -44 | -36 |  | 3.82 |
|  | -22 | -52 | -24 | 141 | 3.86 |
| Occipital cortex | -20 | -84 | -6 | 117 | 3.84 |
|  | 18 | -86 | -4 | 128 | 3.48 |

*Brain activations significant using a $p_{uncorr} < .005$ threshold. Mod = modulation contrast.*

**Supplemental Table S5:**
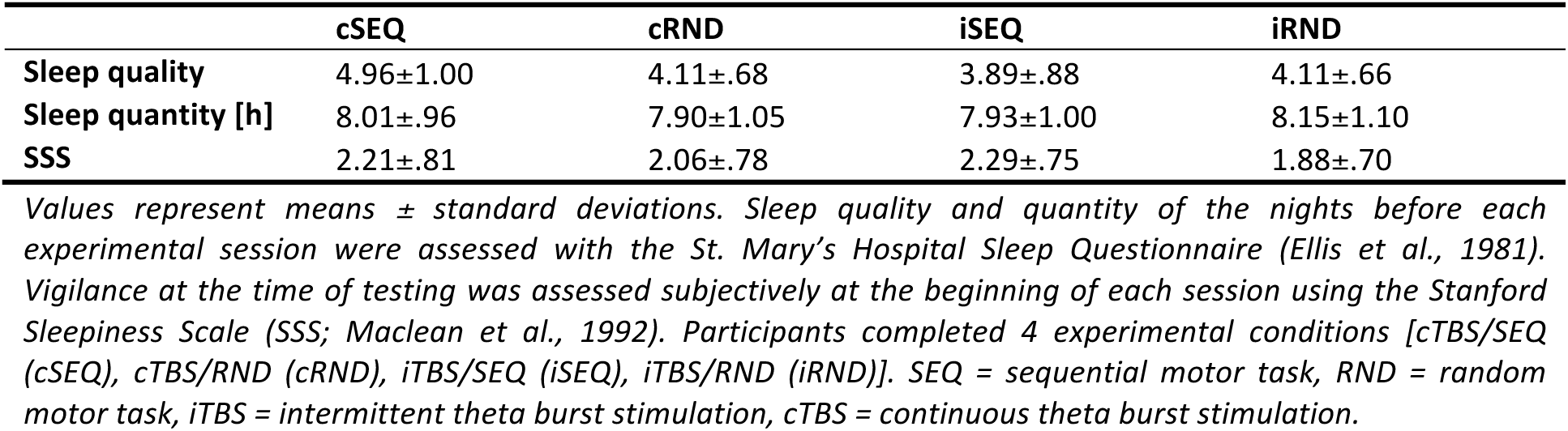
Sleep/vigilance scores within each condition

**Supplemental Table S6:** Individual TMS targets.

| Participant | x mm | y mm | z mm |
| --- | --- | --- | --- |
| P1 | -36 | 16 | 36 |
| P2 | -20 | 30 | 44 |
| P3 | -18 | 18 | 40 |
| P4 | -16 | 18 | 48 |
| P5 | -28 | 16 | 42 |
| P6 | -24 | 16 | 36 |
| P7 | -20 | 20 | 40 |
| P8 | -28 | 18 | 54 |
| P9 | -36 | 16 | 36 |
| P10 | -34 | 16 | 42 |
| P11 | -22 | 26 | 42 |
| P12 | -28 | 24 | 38 |
| P13 | -28 | 32 | 44 |
| P14 | -28 | 18 | 36 |
| P15 | -18 | 24 | 48 |
| P16 | -38 | 12 | 54 |
| P17 | -28 | 12 | 38 |
| P18 | -18 | 14 | 52 |
| P19 | -20 | 12 | 46 |

**Supplemental Table S7:** Data quality metrics for the tissue-corrected GABA+ levels and segmentation results for the DLPFC voxel

| Condition | N | Time | Data quality |  |  | Tissue Fraction |  |  |
| --- | --- | --- | --- | --- | --- | --- | --- | --- |
|  |  |  | GABA SNR | Fit Error [%] | Frequency Offset [ppm] | GM | WM | CSF |
| cSEQ | 18 | pre | 22.89±2.75 | 4.13±1.15 | .0004±.005 | .404±.034 | .529±.039 | .067±.016 |
|  |  | post | 23.67±3.29 | 4.11±1.20 | .0004±.004 | .407±.028 | .515±.043 | .078±.025 |
| cRND | 18 | pre | 22.73±2.97 | 4.29±1.62 | .00009±.003 | .405±.026 | .523±.037 | .072±.022 |
|  |  | post | 22.32±4.305 | 4.20±1.01 | -.0007±.004 | .423±.058 | .505±.070 | .072±.022 |
| iSEQ | 19 | pre | 21.59±3.61 | 4.12±1.93 | .002±.004 | .405±.034 | .524±.045 | .072±.026 |
|  |  | post | 21.68±3.91 | 4.44±1.49 | .001±.003 | .402±.038 | .529±.049 | .069±.023 |
| iRND | 18 | pre | 21.59±3.75 | 4.52±1.87 | .0017±.003 | .403±.022 | .528±.031 | .069±.025 |
|  |  | post | 23.59±3.76 | 4.24±1.04 | .0008±.003 | .400±.026 | .528±.040 | .073±.030 |
| Main effect of time | | | $F_{(1,16)}=.974$<br>$p=.338$ | $F_{(1,16)}=.075$<br>$p=.788$ | $F_{(1,16)}=2.252$<br>$p=.153$ | $F_{(1,16)}=1.039$<br>$p=.323$ | $F_{(1,16)}=1.903$<br>$p=.187$ | $F_{(1,16)}=2.463$<br>$p=.136$ |
| Main effect of stimulation | | | $F_{(1,16)}=1.722$<br>$p=.208$ | $F_{(1,16)}=.323$<br>$p=.577$ | $F_{(1,16)}=22.548$<br>$p=.130$ | $F_{(1,16)}=4.672$<br>**$p=.046$** | $F_{(1,16)}=3.414$<br>$p=.083$ | $F_{(1,16)}=.311$<br>$p=.585$ |
| Main effect of task | | | $F_{(1,16)}=.019$<br>$p=.893$ | $F_{(1,16)}=.188$<br>$p=.670$ | $F_{(1,16)}=.793$<br>$p=.386$ | $F_{(1,16)}=.648$<br>$p=.433$ | $F_{(1,16)}=.407$<br>$p=.532$ | $F_{(1,16)}=.038$<br>$p=.848$ |
| time x stimulation | | | $F_{(1,16)}=.075$<br>$p=.788$ | $F_{(1,16)}=.015$<br>$p=.903$ | $F_{(1,16)}=.394$<br>$p=.539$ | $F_{(1,16)}=2.232$<br>$p=.155$ | $F_{(1,16)}=3.284$<br>$p=.089$ | $F_{(1,16)}=2.433$<br>$p=.138$ |
| time x task | | | $F_{(1,16)}=.364$<br>$p=.555$ | $F_{(1,16)}=.456$<br>$p=.509$ | $F_{(1,16)}=.022$<br>$p=.884$ | $F_{(1,16)}=.698$<br>$p=.416$ | $F_{(1,16)}=.225$<br>$p=.641$ | $F_{(1,16)}=.245$<br>$p=.628$ |
| stimulation x task | | | $F_{(1,16)}=3.709$<br>$p=.072$ | $F_{(1,16)}=.001$<br>$p=.972$ | $F_{(1,16)}=.407$<br>$p=.532$ | $F_{(1,16)}=.288$<br>$p=.599$ | $F_{(1,16)}=.087$<br>$p=.772$ | $F_{(1,16)}=.124$<br>$p=.729$ |
| time x stimulation x task | | | $F_{(1,16)}=1.698$<br>$p=.211$ | $F_{(1,16)}=.311$<br>$p=.585$ | $F_{(1,16)}=.425$<br>$p=.524$ | $F_{(1,16)}=.922$<br>$p=.351$ | $F_{(1,16)}=.005$<br>$p=.943$ | $F_{(1,16)}=5.263$<br>**$p=.036$** |

Values are means ± standard deviations. Statistics are based on a time (2) x stimulation (2) x task (2) repeated measures ANOVA. Participants completed 4 experimental conditions [cTBS/SEQ (cSEQ), cTBS/RND (cRND), iTBS/SEQ (iSEQ), iTBS/RND (iRND)]. GABA = gamma-aminobutyric acid, DLPFC = dorsolateral prefrontal cortex, SNR = signal to noise ratio, FrequencyOffset = average center of the frequency offset, GM = grey matter, WM = white matter, CSF = cerebrospinal fluid, SEQ = sequential motor task, RND = random motor task, iTBS = intermittent theta burst stimulation, cTBS = continuous theta burst stimulation.

**Supplemental Table S8:** Main effect of task practice

| Area | x mm | y mm | z mm | k voxels | T |
| --- | --- | --- | --- | --- | --- |
| Cerebellum | 16 | -56 | -18 | 10715 | 20.92 |
|  | -28 | -54 | -28 |  | 19.27 |
|  | 26 | -50 | -28 |  | 15.68 |
| Motor cortex | -36 | -14 | 62 | 11686 | 16.18 |
|  | 38 | -8 | 60 |  | 14.79 |
|  | 22 | -14 | 50 |  | 14.78 |
| Occipital cortex | -24 | -88 | -18 | 82 | 9.76 |
|  | -28 | -84 | 12 | 194 | 9.50 |
| Frontal cortex | 62 | 10 | 32 | 258 | 9.49 |
|  | 60 | 6 | 24 |  | 9.48 |
| Thalamus/ striatum | 12 | -20 | 18 | 408 | 9.25 |
| Middle temporal cortex | 42 | -66 | 2 | 47 | 8.29 |
| Thalamus | -20 | -14 | 10 | 192 | 8.13 |
|  | -14 | -20 | 12 |  | 7.73 |
| Superior occipital cortex | 20 | -88 | 2 | 19 | 7.70 |
| Middle occipital cortex | -16 | -92 | -6 | 4 | 7.13 |

*Brain activations significant after family-wise error (FWE) correction ( $p_{\text{corr}} < .05$ ) for multiple comparisons over the entire brain volume are reported here.*

## References

1. Albouy, G., Fogel, S., King, B. R., Laventure, S., Benali, H., Karni, A., … Doyon, J. (2015). Maintaining vs. enhancing motor sequence memories: Respective roles of striatal and hippocampal systems. NeuroImage, 108, 423–434. https://doi.org/10.1016/j.neuroimage.2014.12.049

2. Albouy, G., King, B. R., Maquet, P., & Doyon, J. (2013). Hippocampus and striatum: Dynamics and interaction during acquisition and sleep-related motor sequence memory consolidation. Hippocampus, 23(11), 985–1004. https://doi.org/10.1002/hipo.22183

3. Albouy, G., Sterpenich, V., Balteau, E., Vandewalle, G., Desseilles, M., Dang-Vu, T., … Maquet, P. (2008). Both the Hippocampus and Striatum Are Involved in Consolidation of Motor Sequence Memory. Neuron, 58(2), 261–272. https://doi.org/10.1016/j.neuron.2008.02.008

4. Albouy, G., Sterpenich, V., Vandewalle, G., Darsaud, A., Gais, S., Rauchs, G., … Maquet, P. (2012). Neural correlates of performance variability during motor sequence acquisition. NeuroImage, 60(1), 324–331. https://doi.org/10.1016/j.neuroimage.2011.12.049

5. Albouy, G., Sterpenich, V., Vandewalle, G., Darsaud, A., Gais, S., Rauchs, G., … Maquet, P. (2013). Interaction between Hippocampal and Striatal Systems Predicts Subsequent Consolidation of Motor Sequence Memory. PLoS ONE, 8(3), 12–14. https://doi.org/10.1371/journal.pone.0059490

6. Alkhasli, I., Sakreida, K., Mottaghy, F. M., & Binkofski, F. (2019). Modulation of Fronto-Striatal Functional Connectivity Using Transcranial Magnetic Stimulation. Frontiers in Human Neuroscience, 13(June), 1–9. https://doi.org/10.3389/fnhum.2019.00190

7. Ashburner, J., & Friston, K. J. (2005). Unified segmentation. NeuroImage, 26(3), 839– 851. https://doi.org/10.1016/j.neuroimage.2005.02.018

8. Bachtiar, V., Johnstone, A., Berrington, A., Lemke, C., Johansen-Berg, H., Emir, U., & Stagg, C. J. (2018). Modulating regional motor cortical excitability with noninvasive brain stimulation results in neurochemical changes in bilateral motor cortices. Journal of Neuroscience, 38(33), 7327–7336. https://doi.org/10.1523/JNEUROSCI.2853-17.2018

9. Bachtiar, V., Near, J., Johansen-Berg, H., & Stagg, C. J. (2015). Modulation of GABA and resting state functional connectivity by transcranial direct current stimulation. ELife, 4(September 2015), 1–9. https://doi.org/10.7554/eLife.08789

10. Beck, A. T., Epstein, N., Brown, G., & Steer, R. A. (1988). An Inventory for Measuring Clinical Anxiety: Psychometric Properties. Journal of Consulting and Clinical Psychology, 56(6), 893–897. https://doi.org/10.1037/0022-006X.56.6.893

11. Beck, A. T., Ward, C. H., Mendelson, M., Mock, J., & Erbaugh, J. (1961). An Inventory for Measuring Depression. Archives of General Psychiatry, 4(6), 561–571. https://doi.org/10.1001/archpsyc.1961.01710120031004

12. Beynel, L., Appelbaum, L. G., Luber, B., Crowell, C. A., Hilbig, S. A., Lim, W., … Deng, Z. De. (2019). Effects of online repetitive transcranial magnetic stimulation (rTMS) on cognitive processing: A meta-analysis and recommendations for future studies. Neuroscience and Biobehavioral Reviews, 107(August), 47–58. https://doi.org/10.1016/j.neubiorev.2019.08.018

13. Bilek, E., Schafer, A., Ochs, E., Esslinger, C., Zangl, M., Plichta, M. M., … Tost, H. (2013). Application of high-frequency repetitive transcranial magnetic stimulation to the DLPFC alters human prefrontal-hippocampal functional interaction. J Neurosci, 33(16), 7050–7056. https://doi.org/10.1523/jneurosci.3081-12.2013

14. Brainard, D. H. (1997). The Psychophysics Toolbox. Spatial Vision. https://doi.org/10.1163/156856897X00357

15. Burke, M. R., & Coats, R. O. (2016). Dissociation of the rostral and dorsolateral prefrontal cortex during sequence learning in saccades: a TMS investigation. Experimental Brain Research, 234(2), 597–604. https://doi.org/10.1007/s00221-015-4495-2

16. Buysse, D. J., Reynolds, C. F., Monk, T. H., Berman, S. R., & Kupfer, D. J. (1989). The Pittsburgh sleep quality index: A new instrument for psychiatric practice and research. Psychiatry Research, 28(2), 193–213. https://doi.org/10.1016/0165-1781(89)90047-4

17. Chung, S. W., Lewis, B. P., Rogasch, N. C., Saeki, T., Thomson, R. H., Hoy, K. E., … Fitzgerald, P. B. (2017). Demonstration of short-term plasticity in the dorsolateral prefrontal cortex with theta burst stimulation: A TMS-EEG study. Clinical Neurophysiology, 128(7), 1117–1126. https://doi.org/10.1016/j.clinph.2017.04.005

18. Davis, S. W., Luber, B., Murphy, D. L. K., Lisanby, S. H., & Cabeza, R. (2017). Frequency-specific neuromodulation of local and distant connectivity in aging and episodic memory function. Human Brain Mapping, 38(12), 5987–6004. https://doi.org/10.1002/hbm.23803

19. Dayan, E., Herszage, J., Laor-Maayany, R., Sharon, H., & Censor, N. (2018). Neuromodulation of reinforced skill learning reveals the causal function of prefrontal cortex. Human Brain Mapping, (June), 1–9. https://doi.org/10.1002/hbm.24317

20. de Beukelaar, T. T., Van Soom, J., Huber, R., & Wenderoth, N. (2016). A Day Awake Attenuates Motor Learning-Induced Increases in Corticomotor Excitability. Frontiers in Human Neuroscience, 10(August), 138. https://doi.org/10.3389/fnhum.2016.00138

21. Doyon, J., Bellec, P., Amsel, R., Penhune, V., Monchi, O., Carrier, J., … Benali, H. (2009). Contributions of the basal ganglia and functionally related brain structures to motor learning. Behavioural Brain Research, 199(1), 61–75. https://doi.org/10.1016/j.bbr.2008.11.012

22. Doyon, J., Gabitov, E., Vahdat, S., Lungu, O., & Boutin, A. (2018). Current issues related to motor sequence learning in humans. Current Opinion in Behavioral Sciences, 20, 89–97. https://doi.org/10.1016/j.cobeha.2017.11.012

23. Duncan, N. W., Wiebking, C., & Northoff, G. (2014). Associations of regional GABA and glutamate with intrinsic and extrinsic neural activity in humans-A review of multimodal imaging studies. Neuroscience and Biobehavioral Reviews, 47, 36– 52. https://doi.org/10.1016/j.neubiorev.2014.07.016

24. Edden, R. A. E., Oeltzschner, G., Harris, A. D., Puts, N. A. J., Chan, K. L., Boer, V. O., … Barker, P. B. (2016). Prospective frequency correction for macromolecule suppressed GABA editing experiments at 3T. J Magn Reson Imaging, 44(6), 1474–1482. https://doi.org/10.1002/jmri.25304.

25. Edden, R. A. E., Puts, N. A. J., & Barker, P. B. (2012). Macromolecule-suppressed GABA-edited magnetic resonance spectroscopy at 3T. Magnetic Resonance in Medicine, 68(3), 657–661. https://doi.org/10.1002/mrm.24391

26. Edden, R. A. E., Puts, N. A. J., Harris, A. D., Barker, P. B., & Evans, C. J. (2014). Gannet: A batch-processing tool for the quantitative analysis of gamma-aminobutyric acid-edited MR spectroscopy spectra. Journal of Magnetic Resonance Imaging, 40(6), 1445–1452. https://doi.org/10.1002/jmri.24478

27. Ellis, B., Johns, M., Lancaster, R., Raptopoulos, P., Angelopoulos, N., & Priest, R. (1981). The St. Mary’s Hospital Sleep Questionnaire: A Study of Reliability. SLEEP, 4, 93–97. https://doi.org/10.1093/sleep/4.1.93

28. Esslinger, C., Schüler, N., Sauer, C., Gass, D., Mier, D., Braun, U., … Meyer-Lindenberg, A. (2014). Induction and quantification of prefrontal cortical network plasticity using 5 Hz rTMS and fMRI. Human Brain Mapping, 35(1), 140–151. https://doi.org/10.1002/hbm.22165

29. Floyer-Lea, A., Wylezinska, M., Kincses, T., & Matthews, P. M. (2006). Rapid modulation of GABA concentration in human sensorimotor cortex during motor learning. Journal of Neurophysiology, 95(3), 1639–1644. https://doi.org/10.1152/jn.00346.2005

30. Fox, M. D., Buckner, R. L., White, M. P., Greicius, M. D., & Pascual-Leone, A. (2012). Efficacy of transcranial magnetic stimulation targets for depression is related to intrinsic functional connectivity with the subgenual cingulate. Biological Psychiatry, 72(7), 595–603. https://doi.org/10.1016/j.biopsych.2012.04.028

31. Fox, M. D., Halko, M. A., Eldaief, M. C., & Pascual-Leone, A. (2012). Measuring and manipulating brain connectivity with resting state functional connectivity magnetic resonance imaging (fcMRI) and transcranial magnetic stimulation (TMS). NeuroImage, 62(4), 2232–2243. https://doi.org/10.1016/j.neuroimage.2012.03.035

32. Fox, M. D., & Raichle, M. E. (2007). Spontaneous fluctuations in brain activity observed with functional magnetic resonance imaging. Nature Reviews Neuroscience, 8(9), 700–711. https://doi.org/10.1038/nrn2201

33. Fox, M. D., Snyder, A. Z., Vincent, J. L., Corbetta, M., Van Essen, D. C., & Raichle, M. E. (2005). The human brain is intrinsically organized into dynamic, anticorrelated functional networks. Proceedings of the National Academy of Sciences of the United States of America, 102(27), 9673–9678. https://doi.org/10.1073/pnas.0504136102

34. Freedberg, M., Reeves, J. A., Toader, A. C., Hermiller, M. S., Voss, J. L., & Wassermann, E. M. (2019). Persistent enhancement of hippocampal network connectivity by parietal rTMS is reproducible. ENeuro, 6(5), 1–13. https://doi.org/10.1523/ENEURO.0129-19.2019

35. Freedberg, M., Toader, A. C., Wassermann, E. M., & Voss, J. L. (2020). Competitive and cooperative interactions between medial temporal and striatal learning systems. Neuropsychologia, 136(September 2019), 107257. https://doi.org/10.1016/j.neuropsychologia.2019.107257

36. Galea, J. M., Albert, N. B., Ditye, T., & Miall, R. C. (2010). Disruption of the dorsolateral prefrontal cortex facilitates the consolidation of procedural skills. Journal of Cognitive Neuroscience, 22(6), 1158–1164. https://doi.org/10.1162/jocn.2009.21259

37. Gitelman, D. R., Penny, W. D., Ashburner, J., & Friston, K. J. (2003). Modeling regional and psychophysiologic interactions in fMRI: The importance of hemodynamic deconvolution. NeuroImage, 19(1), 200–207. https://doi.org/10.1016/S1053-8119(03)00058-2

38. Gratton, C., Lee, T. G., Nomura, E. M., & D’Esposito, M. (2013). The effect of theta-burst TMS on cognitive control networks measured with resting state fMRI. Frontiers in Systems Neuroscience, 7(December), 1–14. https://doi.org/10.3389/fnsys.2013.00124

39. Gratton, C., Lee, T. G., Nomura, E. M., & D’Esposito, M. (2014). Perfusion MRI indexes variability in the functional brain effects of theta-burst transcranial magnetic stimulation. PLoS ONE, 9(7). https://doi.org/10.1371/journal.pone.0101430

40. Hanlon, C. A., Canterberry, M., Taylor, J. J., DeVries, W., Li, X., Brown, T. R., & George, M. S. (2013). Probing the Frontostriatal Loops Involved in Executive and Limbic Processing via Interleaved TMS and Functional MRI at Two Prefrontal Locations: A Pilot Study. PLoS ONE, 8(7), 1–10. https://doi.org/10.1371/journal.pone.0067917

41. Hanlon, C. A., Dowdle, L. T., Moss, H., Canterberry, M., & George, M. S. (2016). Mobilization of Medial and Lateral Frontal-Striatal Circuits in Cocaine Users and Controls: An Interleaved TMS/BOLD Functional Connectivity Study. Neuropsychopharmacology, 41(13), 3032–3041. https://doi.org/10.1038/npp.2016.114

42. Harris, A. D., Glaubitz, B., Near, J., John Evans, C., Puts, N. A. J., Schmidt-wilcke, T., … Edden, R. A. E. (2014). The Impact of Frequency Drift on GABA-Edited MR Spectroscopy. Magn Reson Med, 72(4), 941–948. https://doi.org/10.1002/mrm.25009

43. Harris, A. D., Puts, N. A. J., & Edden, R. A. E. (2015). Tissue correction for GABA-edited MRS: Considerations of voxel composition, tissue segmentation, and tissue relaxations. Journal of Magnetic Resonance Imaging, 42(5), 1431–1440. https://doi.org/10.1002/jmri.24903

44. Hermans, L., Levin, O., Maes, C., van Ruitenbeek, P., Heise, K. F., Edden, R. A. E., … Cuypers, K. (2018). GABA levels and measures of intracortical and interhemispheric excitability in healthy young and older adults: an MRS-TMS study. Neurobiology of Aging, 65, 168–177. https://doi.org/10.1016/j.neurobiolaging.2018.01.023

45. Hikosaka, O., Nakamura, K., Sakai, K., & Nakahara, H. (2002). Central mechanisms of motor skill learning. Proceedings of the SID, 31(4), 267–365. https://doi.org/10.1016/S0959-4388(02)00307-0

46. Hone-Blanchet, A., Edden, R. A., & Fecteau, S. (2016). Online Effects of Transcranial Direct Current Stimulation in Real Time on Human Prefrontal and Striatal Metabolites. Biol Psychiatry., 80(6), 432–438. https://doi.org/10.1016/j.biopsych.2015.11.008

47. Huang, Y. Z., Edwards, M. J., Rounis, E., Bhatia, K. P., & Rothwell, J. C. (2005). Theta burst stimulation of the human motor cortex. Neuron, 45(2), 201–206. https://doi.org/10.1016/j.neuron.2004.12.033

48. Huang, Y. Z., Rothwell, J. C., Edwards, M. J., & Chen, R. S. (2008). Effect of physiological activity on an NMDA-dependent form of cortical plasticity in human. Cerebral Cortex, 18(3), 563–570. https://doi.org/10.1093/cercor/bhm087

49. Iwabuchi, S. J., Raschke, F., Auer, D. P., Liddle, P. F., Lankappa, S. T., & Palaniyappan, L. (2017). Targeted transcranial theta-burst stimulation alters fronto-insular network and prefrontal GABA. NeuroImage, 146, 395–403. https://doi.org/10.1016/j.neuroimage.2016.09.043

50. King, B. R., Dolfen, N., Gann, M. A., Renard, Z., Swinnen, S. P., & Albouy, G. (2019). Schema and Motor-Memory Consolidation. Psychological Science, 30(7), 963– 978. https://doi.org/10.1177/0956797619847164

51. King, B. R., van Ruitenbeek, P., Leunissen, I., Cuypers, K., Heise, K. F., Santos Monteiro, T., … Swinnen, S. P. (2018). Age-Related Declines in Motor Performance are Associated With Decreased Segregation of Large-Scale Resting State Brain Networks. Cerebral Cortex. https://doi.org/10.1093/cercor/bhx297

52. Kolasinski, J., Hinson, E. L., Divanbeighi Zand, A. P., Rizov, A., Emir, U. E., & Stagg, C. J. (2018). The dynamics of cortical GABA in human motor learning. Journal of Physiology, 1, 271–282. https://doi.org/10.1113/JP276626

53. Lehéricy, S., Benali, H., Van de Moortele, P.-F., Pélégrini-Issac, M., Waechter, T., Ugurbil, K., & Doyon, J. (2005). Distinct basal ganglia territories are engaged in early and advanced motor sequence learning. Proceedings of the National Academy of Sciences of the United States of America, 102(35), 12566–12571. https://doi.org/10.1073/pnas.0502762102

54. Lehéricy, S., Ducros, M., Van De Moortele, P. F., Francois, C., Thivard, L., Poupon, C., … Kim, D. S. (2004). Diffusion Tensor Fiber Tracking Shows Distinct Corticostriatal Circuits in Humans. Annals of Neurology, 55(4), 522–529. https://doi.org/10.1002/ana.20030

55. Levin, O., Weerasekera, A., King, B. R., Heise, K. F., Sima, D. M., Chalavi, S., … Swinnen, S. P. (2019). Sensorimotor cortex neurometabolite levels as correlate of motor performance in normal aging: evidence from a 1H-MRS study. NeuroImage, 202(June), 116050. https://doi.org/10.1016/j.neuroimage.2019.116050

56. Maclean, A. W., Fekken, G. C., Saskin, P., & Knowles, J. B. (1992). Psychometric evaluation of the Stanford Sleepiness Scale. Journal of Sleep Research, 1(1), 35– 39. https://doi.org/10.1111/j.1365-2869.1992.tb00006.x

57. Maes, C., Hermans, L., Pauwels, L., Chalavi, S., Leunissen, I., Levin, O., … Swinnen, S. P. (2018). Age-related differences in GABA levels are driven by bulk tissue changes. Human Brain Mapping, 39(9), 3652–3662. https://doi.org/10.1002/hbm.24201

58. Marjańska, M., Lehéricy, S., Valabregue, R., Popa, T., Bonnet, C., Gallea, C., & Coudert, M. (2013). Brain dynamic neurochemical changes in dystonic patients: a magnetic resonance spectroscopy study. Mov Disord., 28(2), 201–209. https://doi.org/10.1002/mds.25279

59. Mastropasqua, C., Bozzali, M., Ponzo, V., Giulietti, G., Caltagirone, C., Cercignani, M., & Koch, G. (2014). Network based statistical analysis detects changes induced by continuous theta-burst stimulation on brain activity at rest. Frontiers in Psychiatry, 5(AUG), 1–7. https://doi.org/10.3389/fpsyt.2014.00097

60. Mescher, M., Merkle, H., Kirsch, J., Garwood, M., & Gruetter, R. (1998). Simultaneous in vivo spectral editing and water suppression. NMR in Biomedicine, 11(6), 266–272. https://doi.org/10.1002/(SICI)1099-1492(199810)11:6<266::AID-NBM530>3.0.CO;2-J

61. Mikkelsen, M., Barker, P. B., Bhattacharyya, P. K., Brix, M. K., Buur, P. F., Cecil, K. M., … Edden, R. A. E. (2017). Big GABA: Edited MR spectroscopy at 24 research sites. NeuroImage, 159, 32–45. https://doi.org/10.1016/j.neuroimage.2017.07.021

62. Mikkelsen, M., Loo, R. S., Puts, N. A. J., Edden, R. A. E., & Harris, A. D. (2018). Designing GABA-edited magnetic resonance spectroscopy studies: Considerations of scan duration, signal-to-noise ratio and sample size. Journal of Neuroscience Methods. https://doi.org/10.1016/j.jneumeth.2018.02.012

63. Mikkelsen, M., Rimbault, D. L., Barker, P. B., Bhattacharyya, P. K., Brix, M. K., Buur, P. F., … Edden, R. A. E. (2019). Big GABA II: Water-referenced edited MR spectroscopy at 25 research sites. NeuroImage, 191, 537–548. https://doi.org/10.1016/j.neuroimage.2019.02.059

64. Monteiro, T. S., Zivari Adab, H., Chalavi, S., Gooijers, J., King, B. (Bradley) R., Cuypers, K., … Swinnen, S. P. (2020). Reduced Modulation of Task-Related Connectivity Mediates Age-Related Declines in Bimanual Performance. Cerebral Cortex, (January 2001), 1–15. https://doi.org/10.1093/cercor/bhaa021

65. Mullins, P. G., McGonigle, D. J., O’Gorman, R. L., Puts, N. A. J., Vidyasagar, R., Evans, C. J., … Edden, R. A. E. (2014). Current practice in the use of MEGA-PRESS spectroscopy for the detection of GABA. Neuroimage., 86, 43–52. https://doi.org/10.1016/j.neuroimage.2012.12.004

66. Muto, V., Jaspar, M., Meyer, C., Kussé, C., Chellappa, S. L., Degueldre, C., … Maquet, P. (2016). Local modulation of human brain responses by circadian rhythmicity and sleep debt. Science, 353(6300), 687 LP – 690. https://doi.org/10.1126/science.aad2993

67. Near, J., Edden, R., Evans, C. J., Paquin, R., Harris, A., & Jezzard, P. (2015). Frequency and phase drift correction of magnetic resonance spectroscopy data by spectral registration in the time domain. Magnetic Resonance in Medicine, 73(1), 44–50. https://doi.org/10.1002/mrm.25094

68. Nissen, M. J., & Bullemer, P. (1987). Attentional requirements of learning: Evidence from performance measures. Cognitive Psychology, 19(1), 1–32. https://doi.org/10.1016/0010-0285(87)90002-8

69. Oldfield, R. C. (1971). The assessment and analysis of handedness: The Edinburgh inventory. Neuropsychologia, 9(1), 97–113. https://doi.org/10.1016/0028-3932(71)90067-4

70. Ott, D. V. M., Ullsperger, M., Jocham, G., Neumann, J., & Klein, T. A. (2011). Continuous theta-burst stimulation (cTBS) over the lateral prefrontal cortex alters reinforcement learning bias. NeuroImage, 57(2), 617–623. https://doi.org/10.1016/j.neuroimage.2011.04.038

71. Pascual-Leone, A., Wassermann, E. M., Grafman, J., & Hallett, M. (1996). The role of the dorsolateral prefrontal cortex in implicit procedural learning. Experimental Brain Research, 107(3), 479–485. https://doi.org/10.1007/BF00230427

72. Penhune, V. B., & Steele, C. J. (2012). Parallel contributions of cerebellar, striatal and M1 mechanisms to motor sequence learning. Behavioural Brain Research, 226(2), 579–591. https://doi.org/10.1016/j.bbr.2011.09.044

73. Puts, N. A. J., & Edden, R. A. E. (2012). In vivo magnetic resonance spectroscopy of GABA: A methodological review. Progress in Nuclear Magnetic Resonance Spectroscopy, 60, 29–41. https://doi.org/10.1016/j.pnmrs.2011.06.001

74. Robertson, E. M., Tormos, J. M., Maeda, F., & Pascual-Leone, A. (2001). The role of the dorsolateral prefrontal cortex during sequence learning is specific for spatial information. Cerebral Cortex, 11(7), 628–635. https://doi.org/10.1093/cercor/11.7.628

75. Rothman, D. L., Petroff, O. A. C., Behar, K. L., & Mattson, R. H. (1993). Localized 1H NMR measurements of γ-aminobutyric acid in human brain in vivo. Proceedings of the National Academy of Sciences of the United States of America, 90(12), 5662–5666. https://doi.org/10.1073/pnas.90.12.5662

76. Rounis, E., Stephan, K. E., Lee, L., Siebner, H. R., Pesenti, A., Friston, K. J., … Frackowiak, R. S. J. (2006). Acute changes in frontoparietal activity after repetitive transcranial magnetic stimulation over the dorsolateral prefrontal cortex in a cued reaction time task. Journal of Neuroscience, 26(38), 9629–9638. https://doi.org/10.1523/JNEUROSCI.2657-06.2006

77. Sack, A. T., Kadosh, R. C., Schuhmann, T., Moerel, M., Walsh, V., & Goebel, R. (2009). Optimizing Functional Accuracy of TMS in Cognitive Studies: A Comparison of Methods. Journal of Cognitive Neuroscience, 21(2), 207–221. https://doi.org/10.1162/jocn.2009.21126

78. Sampaio-Baptista, C., Filippini, N., Stagg, C. J., Near, J., Scholz, J., & Johansen-Berg, H. (2015). Changes in functional connectivity and GABA levels with long-term motor learning. NeuroImage, 106, 15–20. https://doi.org/10.1016/j.neuroimage.2014.11.032

79. Shang, Y., Chang, D., Zhang, J., Peng, W., Song, D., Gao, X., & Wang, Z. (2019). Theta-burst transcranial magnetic stimulation induced functional connectivity changes between dorsolateral prefrontal cortex and default-mode-network. Brain Imaging and Behavior. https://doi.org/10.1007/s11682-019-00139-y

80. Smarr, B. L., Jennings, K. J., Driscoll, J. R., & Kriegsfeld, L. J. (2014). A Time to Remember: The Role of Circadian Clocks in Learning and Memory. Curr. Opin. Biotechnol., 29(3), 146–155. https://doi.org/10.1037/a0035963

81. Stagg, C. J. (2014). Magnetic Resonance Spectroscopy as a tool to study the role of GABA in motor-cortical plasticity. NeuroImage, 86, 19–27. https://doi.org/10.1016/j.neuroimage.2013.01.009

82. Stagg, C. J., Bachtiar, V., & Johansen-Berg, H. (2011). The role of GABA in human motor learning. Current Biology, 21(6), 480–484. https://doi.org/10.1016/j.cub.2011.01.069

83. Stagg, C. J., Best, J. G., Stephenson, M. C., O’Shea, J., Wylezinska, M., Kineses, Z. T., … Johansen-Berg, H. (2009). Polarity-sensitive modulation of cortical neurotransmitters by transcranial stimulation. Journal of Neuroscience, 29(16), 5202–5206. https://doi.org/10.1523/JNEUROSCI.4432-08.2009

84. Stagg, C. J., Wylezinska, M., Matthews, P. M., Jezzard, P., Rothwell, J. C., & Bestmann, S. (2009). Neurochemical Effects of Theta Burst Stimulation as Assessed by Magnetic Resonance Spectroscopy. Journal of Neurophysiology, 101(Oldfield 1971), 2872–2877. https://doi.org/10.1152/jn.91060.2008.

85. Steele, C. J., & Penhune, V. B. (2010). Specific increases within global decreases: a functional magnetic resonance imaging investigation of five days of motor sequence learning. J Neurosci, 30(24), 8332–8341. https://doi.org/10.1523/jneurosci.5569-09.2010

86. Tambini, A., Nee, D. E., & D’Esposito, M. (2018). Hippocampal-targeted Theta-burst Stimulation Enhances Associative Memory Formation. Journal of Cognitive Neuroscience, 30(10), 1452–1472. https://doi.org/10.1162/jocn_a_01300

87. Tang, Y., Jiao, X., Wang, J., Zhu, T., Zhou, J., Qian, Z., … Wang, J. (2019). Dynamic Functional Connectivity Within the Fronto-Limbic Network Induced by Intermittent Theta-Burst Stimulation: A Pilot Study. Frontiers in Neuroscience, 13(September), 1–9. https://doi.org/10.3389/fnins.2019.00944

88. Toro, R., Fox, P. T., & Paus, T. (2008). Functional coactivation map of the human brain. Cerebral Cortex, 18(11), 2553–2559. https://doi.org/10.1093/cercor/bhn014

89. Tunovic, S., Press, D. Z., & Robertson, E. M. (2014). A Physiological Signal That Prevents Motor Skill Improvements during Consolidation. Journal of Neuroscience, 34(15), 5302–5310. https://doi.org/10.1523/JNEUROSCI.3497-13.2014

90. Tzourio-Mazoyer, N., Landeau, B., Papathanassiou, D., Crivello, F., Etard, O., Delcroix, N., … Joliot, M. (2002). Automated anatomical labeling of activations in SPM using a macroscopic anatomical parcellation of the MNI MRI single-subject brain. NeuroImage, 15(1), 273–289. https://doi.org/10.1006/nimg.2001.0978

91. van der Werf, Y. D., Sanz-Arigita, E. J., Menning, S., & van den Heuvel, O. a. (2010). Modulating spontaneous brain activity using repetitive transcranial magnetic stimulation. BMC Neuroscience, 11(1), 145. https://doi.org/10.1186/1471-2202-11-145

92. Van Holstein, M., Froböse, M. I., O’Shea, J., Aarts, E., & Cools, R. (2018). Controlling striatal function via anterior frontal cortex stimulation. Scientific Reports, 8(1), 1–13. https://doi.org/10.1038/s41598-018-21346-5

93. van Polanen, V., Rens, G., & Davare, M. (2019). The Role of the Anterior Intraparietal Sulcus and the Lateral Occipital Cortex in Fingertip Force Scaling and Weight Perception During Object Lifting. *BioRxiv*. https://doi.org/10.1101/2019.12.20.883918

94. Wang, J. X., Rogers, L. M., Gross, E. Z., Ryals, A. J., Dokucu, M. E., Brandstatt, K. L., … Voss, J. L. (2014). Targeted enhancement of cortical-hippocampal brain networks and associative memory. Science, 345(6200), 1054–1057. https://doi.org/10.1126/science.1252900

95. Xue, S. W., Guo, Y., Peng, W., Zhang, J., Chang, D., Zang, Y. F., & Wang, Z. (2017). Increased low-frequency resting-state brain activity by high-frequency repetitive TMS on the left dorsolateral prefrontal cortex. Frontiers in Psychology, 8(DEC), 1–8. https://doi.org/10.3389/fpsyg.2017.02266

96. Zivari Adab, H., Chalavi, S., Monteiro, T. S., Gooijers, J., Dhollander, T., Mantini, D., & Swinnen, S. P. (2020). Fiber-specific variations in anterior transcallosal white matter structure contribute to age-related differences in motor performance. NeuroImage, 209(January), 116530. https://doi.org/10.1016/j.neuroimage.2020.116530

## Supplemental References

2. Albouy, G., Sterpenich, V., Balteau, E., Vandewalle, G., Desseilles, M., Dang-Vu, T., … Maquet, P. (2008). Both the Hippocampus and Striatum Are Involved in Consolidation of Motor Sequence Memory. Neuron, 58(2), 261–272. https://doi.org/10.1016/j.neuron.2008.02.008

3. Albouy, G., Sterpenich, V., Vandewalle, G., Darsaud, A., Gais, S., Rauchs, G., … Maquet, P. (2013). Interaction between Hippocampal and Striatal Systems Predicts Subsequent Consolidation of Motor Sequence Memory. PLoS ONE, 8(3), 12–14. https://doi.org/10.1371/journal.pone.0059490

4. Alkhasli, I., Sakreida, K., Mottaghy, F. M., & Binkofski, F. (2019). Modulation of Fronto-Striatal Functional Connectivity Using Transcranial Magnetic Stimulation. Frontiers in Human Neuroscience, 13(June), 1–9. https://doi.org/10.3389/fnhum.2019.00190

5. Amiez, C., & Petrides, M. (2014). Neuroimaging evidence of the anatomo-functional organization of the human cingulate motor areas. Cerebral Cortex, 24(3), 563–578. https://doi.org/10.1093/cercor/bhs329

6. Bilek, E., Schafer, A., Ochs, E., Esslinger, C., Zangl, M., Plichta, M. M., … Tost, H. (2013). Application of high-frequency repetitive transcranial magnetic stimulation to the DLPFC alters human prefrontal-hippocampal functional interaction. J Neurosci, 33(16), 7050– 7056. https://doi.org/10.1523/jneurosci.3081-12.2013

7. Bischoff-Grethe, A., Goedert, K. M., Willingham, D. T., & Grafton, S. T. (2004). Neural Substrates of Response-based Sequence Learning using fMRI. Journal of Cognitive Neuroscience, 16(1), 127–138. https://doi.org/10.1162/089892904322755610

8. Cao, N., Pi, Y., Liu, K., Meng, H., Wang, Y., Zhang, J., … Tan, X. (2018). Inhibitory and facilitatory connections from dorsolateral prefrontal to primary motor cortex in healthy humans at rest—An rTMS study. Neuroscience Letters, 687(June), 82–87. https://doi.org/10.1016/j.neulet.2018.09.032

9. Chung, S. W., Lewis, B. P., Rogasch, N. C., Saeki, T., Thomson, R. H., Hoy, K. E., … Fitzgerald, P. B. (2017). Demonstration of short-term plasticity in the dorsolateral prefrontal cortex with theta burst stimulation: A TMS-EEG study. Clinical Neurophysiology, 128(7), 1117–1126. https://doi.org/10.1016/j.clinph.2017.04.005

10. Civardi, C., Cantello, R., Asselman, P., & Rothwell, J. C. (2001). Transcranial Magnetic Stimulation Can Be Used to Test Connections to Primary Motor Areas from Frontal and Medial Cortex in Humans. NeuroImage, 14(6), 1444–1453. https://doi.org/10.1006/nimg.2001.0918

11. Dayan, E., Herszage, J., Laor-Maayany, R., Sharon, H., & Censor, N. (2018). Neuromodulation of reinforced skill learning reveals the causal function of prefrontal cortex. Human Brain Mapping, (June), 1–9. https://doi.org/10.1002/hbm.24317

12. Do, M., Kirkovski, M., Davies, C. B., Bekkali, S., Byrne, L. K., & Enticott, P. G. (2018). Intra- and inter-regional priming of ipsilateral human primary motor cortex with continuous theta burst stimulation does not induce consistent neuroplastic effects. Frontiers in Human Neuroscience, 12(March), 1–8. https://doi.org/10.3389/fnhum.2018.00123

13. Ellis, B., Johns, M., Lancaster, R., Raptopoulos, P., Angelopoulos, N., & Priest, R. (1981). The St. Mary’s Hospital Sleep Questionnaire: A Study of Reliability. SLEEP, 4, 93–97. https://doi.org/10.1093/sleep/4.1.93

14. Fernández-Seara, M. A., Aznárez-Sanado, M., Mengual, E., Loayza, F. R., & Pastor, M. A. (2009). Continuous performance of a novel motor sequence leads to highly correlated striatal and hippocampal perfusion increases. NeuroImage, 47(4), 1797–1808. https://doi.org/10.1016/j.neuroimage.2009.05.061

15. Fierro, B., De Tommaso, M., Giglia, F., Giglia, G., Palermo, A., & Brighina, F. (2010). Repetitive transcranial magnetic stimulation (rTMS) of the dorsolateral prefrontal cortex (DLPFC) during capsaicin-induced pain: Modulatory effects on motor cortex excitability. Experimental Brain Research, 203(1), 31–38. https://doi.org/10.1007/s00221-010-2206-6

16. Fischer, S. (2005). Motor Memory Consolidation in Sleep Shapes More Effective Neuronal Representations. Journal of Neuroscience, 25(49), 11248–11255. https://doi.org/10.1523/JNEUROSCI.1743-05.2005

17. Galea, J. M., Albert, N. B., Ditye, T., & Miall, R. C. (2010). Disruption of the dorsolateral prefrontal cortex facilitates the consolidation of procedural skills. Journal of Cognitive Neuroscience, 22(6), 1158–1164. https://doi.org/10.1162/jocn.2009.21259

18. Gheysen, F., Lasne, G., Pélégrini-Issac, M., Albouy, G., Meunier, S., Benali, H., … Popa, T. (2016). Taking the brakes off the learning curve. Human Brain Mapping, 1691(August 2016), 1676–1691. https://doi.org/10.1002/hbm.23489

19. Grafton, S. T., Hazeltine, E., & Ivry, R. (1995). Functional mapping of sequence learning in normal humans. Journal of Cognitive Neuroscience, 7(4), 497–510. https://doi.org/10.1162/jocn.1995.7.4.497

20. Grafton, S. T., Hazeltine, E., & Ivry, R. B. (1998). Abstract and effector-specific representations of motor sequences identified with pet. Journal of Neuroscience, 18(22), 9420–9428. https://doi.org/10.1523/jneurosci.18-22-09420.1998

21. Hone-Blanchet, A., Edden, R. A., & Fecteau, S. (2016). Online Effects of Transcranial Direct Current Stimulation in Real Time on Human Prefrontal and Striatal Metabolites. Biol Psychiatry., 80(6), 432–438. https://doi.org/10.1016/j.biopsych.2015.11.008

22. Lungu, O., Monchi, O., Albouy, G., Jubault, T., Ballarin, E., Burnod, Y., & Doyon, J. (2014). Striatal and hippocampal involvement in motor sequence chunking depends on the learning strategy. PLoS ONE, 9(8), 25–27. https://doi.org/10.1371/journal.pone.0103885

23. Maclean, A. W., Fekken, G. C., Saskin, P., & Knowles, J. B. (1992). Psychometric evaluation of the Stanford Sleepiness Scale. Journal of Sleep Research, 1(1), 35–39. https://doi.org/10.1111/j.1365-2869.1992.tb00006.x

24. Mikkelsen, M., Barker, P. B., Bhattacharyya, P. K., Brix, M. K., Buur, P. F., Cecil, K. M., … Edden, R. A. E. (2017). Big GABA: Edited MR spectroscopy at 24 research sites. NeuroImage, 159, 32–45. https://doi.org/10.1016/j.neuroimage.2017.07.021

25. Mikkelsen, M., Loo, R. S., Puts, N. A. J., Edden, R. A. E., & Harris, A. D. (2018). Designing GABA-edited magnetic resonance spectroscopy studies: Considerations of scan duration, signal-to-noise ratio and sample size. Journal of Neuroscience Methods. https://doi.org/10.1016/j.jneumeth.2018.02.012

26. Mikkelsen, M., Rimbault, D. L., Barker, P. B., Bhattacharyya, P. K., Brix, M. K., Buur, P. F., … Edden, R. A. E. (2019). Big GABA II: Water-referenced edited MR spectroscopy at 25 research sites. NeuroImage, 191, 537–548. https://doi.org/10.1016/j.neuroimage.2019.02.059

27. Oishi, K., Toma, K., Bagarinao, E. T., Matsuo, K., Nakai, T., Chihara, K., & Fukuyama, H. (2005). Activation of the precuneus is related to reduced reaction time in serial reaction time tasks. Neuroscience Research, 52(1), 37–45. https://doi.org/10.1016/j.neures.2005.01.008

28. Penhune, V. B., & Doyon, J. (2002). Dynamic cortical and subcortical networks in learning and delayed recall of timed motor sequences. Journal of Neuroscience, 22(4), 1397– 1406. https://doi.org/10.1523/jneurosci.22-04-01397.2002

29. Penhune, V. B., & Doyon, J. (2005). Cerebellum and M1 interaction during early learning of timed motor sequences. NeuroImage, 26(3), 801–812. https://doi.org/10.1016/j.neuroimage.2005.02.041

30. Rens, G., van Polanen, V., Botta, A., Gann, M. A., Orban de Xivry, J.-J., & Davare, M. (2020). Sensorimotor expectations bias motor resonance during observation of object lifting: The causal role of pSTS. *The Journal of Neuroscience*, JN-RM-2672–19. https://doi.org/10.1523/JNEUROSCI.2672-19.2020

31. Robertson, E. M., Tormos, J. M., Maeda, F., & Pascual-Leone, A. (2001). The role of the dorsolateral prefrontal cortex during sequence learning is specific for spatial information. Cerebral Cortex, 11(7), 628–635. https://doi.org/10.1093/cercor/11.7.628

32. Rollnik, J. D., Schubert, M., & Dengler, R. (2000). Subthreshold prefrontal repetitive transcranial magnetic stimulation reduces motor cortex excitability. Muscle & Nerve, 23(1), 112–114. https://doi.org/10.1002/(SICI)1097-4598(200001)23:1<112::AID-MUS15>3.0.CO;2-B

33. Sakai, K., Ramnani, N., & Passingham, R. E. (2002). Learning of sequences of finger movements and timing: Frontal lobe and action-oriented representation. Journal of Neurophysiology, 88(4), 2035–2046. https://doi.org/10.1152/jn.2002.88.4.2035

34. Schendan, H. E., Searl, M. M., Melrose, R. J., & Stern, C. E. (2003). An fMRI study of the role of the medial temporal lobe in implicit and explicit sequence learning. Neuron, 37(6), 1013–1025. https://doi.org/10.1016/S0896-6273(03)00123-5

35. Sterpenich, V., Albouy, G., Boly, M., Vandewalle, G., Darsaud, A., Balteau, E., … Maquet, P. (2007). Sleep-related hippocampo-cortical interplay during emotional memory recollection. PLoS Biology, 5(11), 2709–2722. https://doi.org/10.1371/journal.pbio.0050282

